# Temperature alone is not enough: food-web context determines evolutionary responses to warming

**DOI:** 10.1101/2024.05.06.592770

**Authors:** Ze-Yi Han, Yaning Yuan, Katrina DeWitt, Andrea Yammine, Daniel J. Wieczynski, Masayuki Onishi, Jean P. Gibert

**Author notes:** To whom correspondence should be addressed:* Ze-Yi Han, Department of Biology, Duke University, Durham, NC, USA. Statement of authorship:* ZH and JPG conceived the study; ZH and JPG designed the experimental work. ZH collected the data with support from AY, KD and YY. ZH analyzed the data with support from DJW and JPG; ZH wrote the first version of the manuscript; all authors contributed significantly to subsequent versions. Data accessibility statement:* All annotated code and data are available at our dedicated repository (https://github.com/ZeYiHan/Temp_Pred_Evo).

## Abstract

Global warming is reshaping food webs globally. Rapid evolution has been proposed as a buffer against climate change, but how simultaneous shifts in biotic and abiotic environments may influence evolution is unknown. Using experimental evolution and mathematical modeling in microbial food webs of prey algae and ciliate predators, we tested 1) how temperature affects prey evolution and 2) how the food-web context––i.e., predator identity, abundance, and competition among predators–– mediates prey evolutionary dynamics. We found that temperature alone does not drive prey evolution unless predators are present, and food-web context determines ensuing evolutionary dynamics. These seemingly complex evolutionary responses are predictable from the joint effects of temperature-dependent, predator-specific predation rates, and the emergence of temperature-dependent prey plasticity. We reveal that evolutionary outcomes under warming are shaped by the broader food web context of species, suggesting that the same species may exhibit different eco-evolutionary responses in different food webs under novel climates.

**SIGNIFICANCE:** Predicting how species evolve under climate change is critical for understanding future changes in food webs. Evolutionary responses have long been known to be driven by environmental change—like temperature—but whether and how ecological interactions influence this process is unknown. Using experimental evolution and mathematical modeling, we show that temperature alone does not drive prey evolution. Instead, the broader food webs context—predator identity, abundance, and competition—mediates how species evolve under warming. Additionally, we demonstrate that prey evolution depends on temperature-dependent predator-specific predation rates and prey plasticity. Our findings highlight that the same species may evolve differently within different food webs, urging the need to integrate ecological interactions when forecasting evolutionary responses to climate change.

## INTRODUCTION

Global warming is reshaping food webs worldwide (Barbour & Gibert 2021) through changes in population growth (Dell *et al*. 2011; Frazier *et al*. 2006; Kontopoulos *et al*. 2020) and species interactions (Blois *et al*. 2013; Dell *et al*. 2014). Additionally, increasing temperature raises metabolic costs (Clarke 2006; Clarke & Fraser 2004), forcing predators to consume more prey (Sheridan & Bickford 2011) while gaining diminishing energetic returns (Barneche *et al*. 2021). This reduced energetic intake results in the energetic choking of upper food-web trophic levels, food web rewiring (Barbour & Gibert 2021; Bartley *et al*. 2019), compositional turnover (Komatsu *et al*. 2019), and trophic collapse (Ullah *et al*. 2018; Voigt *et al*. 2003; Zarnetske *et al*. 2012).

Species evolution both influences and is influenced by novel biotic and abiotic conditions. For example, rising temperature could directly influence evolution through effects on organismal metabolism (Alton *et al*. 2024; Clarke 2003), morphology (Diamond *et al*. 2017; Yampolsky *et al*. 2014), and fitness (van Heerwaarden & Sgrò 2021; Padfield *et al*. 2016). On the other hand, predation and competition within food webs can influence evolution directly (Abrams 2000; Frickel *et al*. 2017), for example, by selecting in favor of better defended prey that are worse at competing for resources (Yoshida *et al*. 2004). In a warming world, however, changes to abiotic drivers of selection, like temperature, can influence biotic drivers of selection, like predation or competition, urging joint study of these effects.

Indeed, temperature influences demographic and functional traits asymmetrically among interacting species (Dell *et al*. 2014; Gibert *et al*. 2022), shifting species interactions and the potential for species rapid evolution under warming (Kordas *et al*. 2011; Tseng & O’Connor 2015). Higher temperature and predation can also trigger plastic responses that fundamentally alter ecological dynamics, species interactions, and ensuing selection (Agrawal 2001). Species evolutionary responses in a warming world are thus likely jointly determined by the effects that rising temperature and resulting shifts in species interactions (i.e., food web context) may have on phenotypic plasticity and selection. However, how these joint effects may affect evolutionary responses within food webs in a warming world is not known, thus greatly hindering our ability to anticipate the fate of species in new climates.

We addressed this central issue through a combination of mathematical models and microcosm experiments in a model system of global distribution and importance (Bar-On *et al*. 2018; Herron *et al*. 2019): the interaction between the unicellular green alga *Chlamydomonas reinhardtii* and its ciliate predators, which co-occur within broader microbial food webs across ecosystems, from soils to wetlands (Foissner *et al*. 2009). While evolutionary change operates in all food webs, addressing how it might change in novel climates is logistically impossible to test in nature. Our microbial system breaks down this intractability issue while remaining relevant to the natural world through the pivotal role of microbes in all ecosystems worldwide (Foissner *et al*. 2009). We thus assembled tractable microcosm food webs and studied the combined effects of temperature, predation, and predator competition (as a proxy for food web context), on prey evolution. We tracked population dynamics, rapid shifts in prey genetic makeup (i.e., rapid evolution), and plastic phenotypic changes, to disentangle how temperature and food web context co-determine eco-evolutionary outcomes within these microbial food webs across three temperatures.

## RESULTS

### Predation maintains genetic diversity and influences eco-evolutionary outcomes

To make testable predictions on how predation could influence prey evolution, we used a mathematical model of rapid evolution that keeps track of two genetically distinct prey strains under predation (see Methods). The model suggests that different prey strains cannot coexist in the absence of predators –classically, unless intra-strain competition is higher than inter-strain competition– (Appendix I Equation 1-2). Predation, however, facilitates the invasion of the strain that would otherwise be lost (Appendix I Equations 1,3), thus maintaining genetic diversity. We tested these theoretical predictions using an experimental predator-prey microbial system where the alga *Chlamydomonas reinhardtii* was preyed upon by one of three possible ciliate protist species (*Glaucoma sp.*, *Tetrahymena pyriformis*, *Paramecium caudatum*, see Methods). The algal prey population was composed of two genetically and phenotypically distinct strains – i.e., the fluorescently tagged wild type (WT) and untagged *vfl1-1* (*v*ariable *fl*agella) genotypes (Fig. 1). We kept track of prey abundances, phenotypes, and genetic frequencies over time to quantify both plastic and evolutionary responses (see Methods). Our empirical results corroborated our model predictions. In the absence of predators, WT consistently outcompeted *vfl1-1*; However, predation either 1) facilitated the persistence of *vfl1-1* that was lost under control conditions, or 2) slowed down the rate of *vfl1-1* loss (Fig. 2a-d; Appendix II Figure 1-2). By altering the temporal dynamics of prey genetic frequencies, predator-prey interactions can determine evolutionary outcomes.

**Figure 1.**
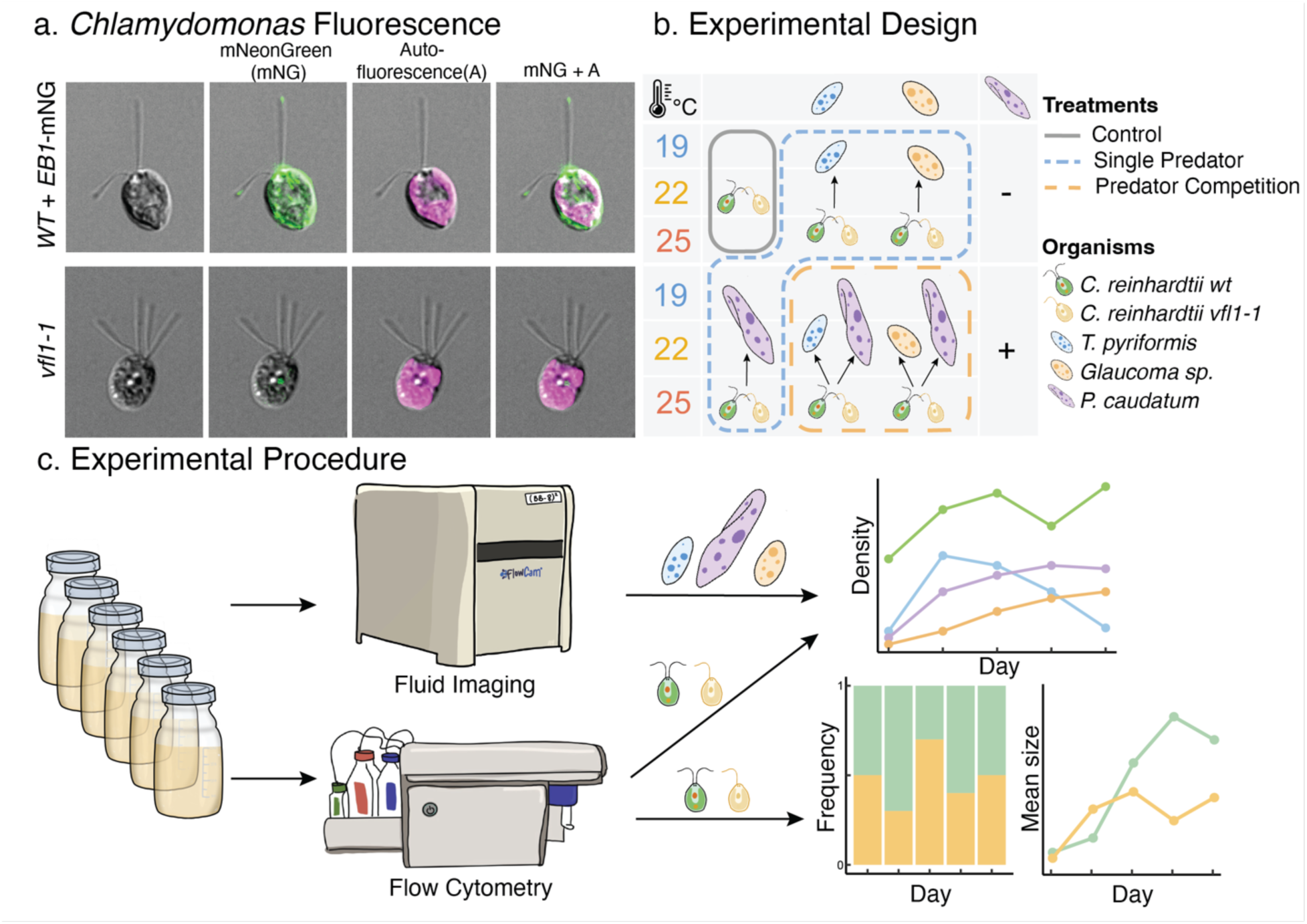
Experimental organisms and designs. a) Images of a *C. reinhardtii* wild-type strain expressing mNeonGreen (mNG)-tagged EB1 protein and a *vfl1-1* mutant strain. *vfl1-1* cells have variable numbers of flagella ranging between zero to more than four. b) Factorial design of the experiment, showing 3 temperatures by 3 predations by 2 competition treatments. c) Experimental procedures of sampling methods.

**Figure 2.**
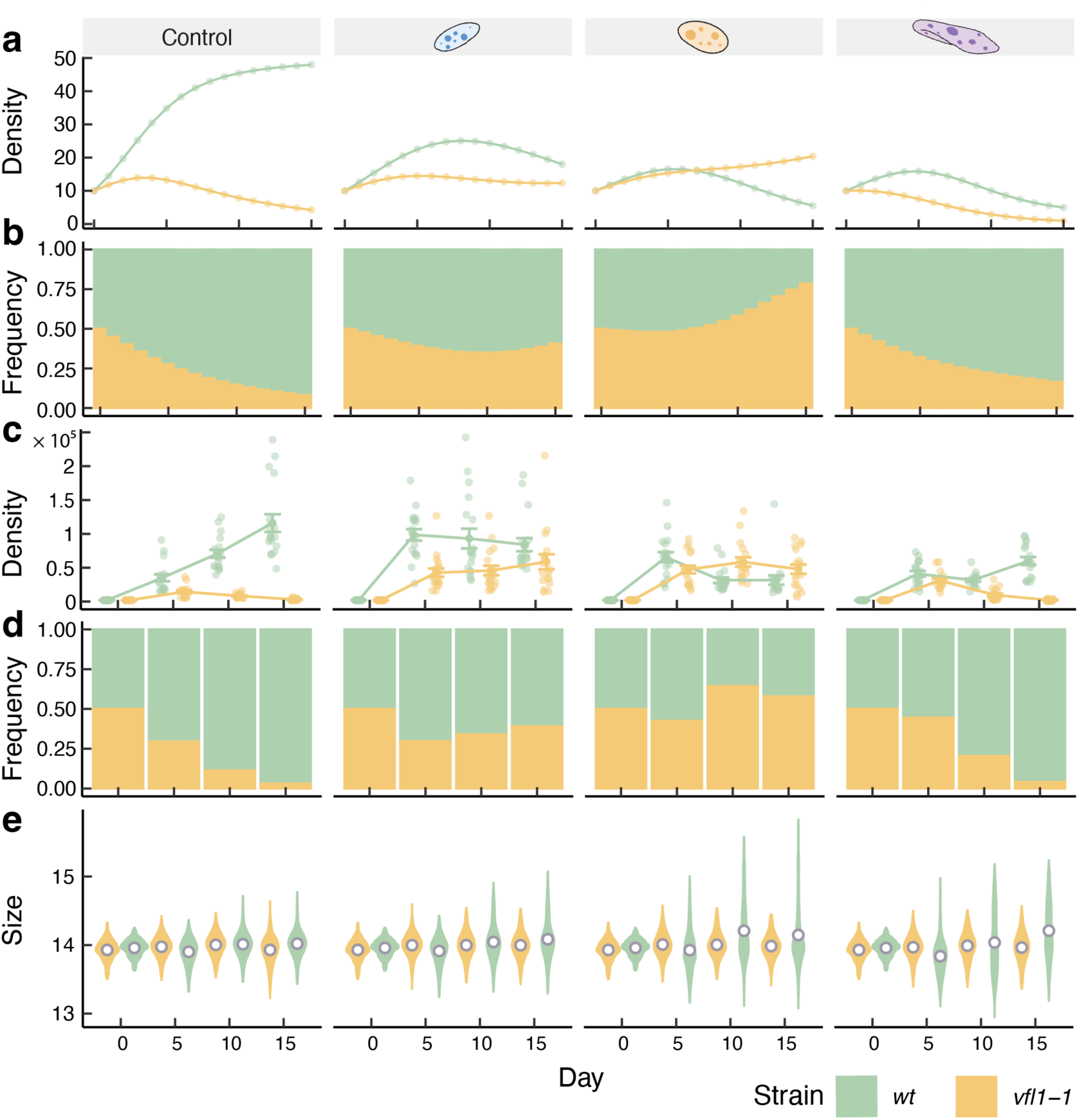
Model prediction and empirical data of prey clonal dynamics and prey body size. *C. reinhardtii* population density and genetic frequencies predicted by mathematical models (a-b; Specific parameter see Appendix I Table 1) and from empirical data (c-d). Panel e shows the size of *C. reinhardtii* particles over time. Yellow represents *vfl1-1* mutant strain and green represents WT.

Different predators selected for WT and *vfl1-1* differently, as indicated by differences in *vfl1-1* frequencies across predator treatments (Fig. 2a-d; Appendix IV Table 1-2). Interestingly, our model can reproduce these effects and suggest that differences in predator attack rates across genotypes alone should be sufficient to explain differences in prey evolutionary outcomes across predators (Fig. 2a-b, model parameter see Appendix I Table 1). This mathematical understanding thus offers a simple explanation for otherwise seemingly complex dynamics, and shows that predator-specific differences in predation rates influence the eco-evolutionary process and prey genetic makeup.

### Prey traits influence eco-evolutionary outcomes

In our experimental system, predation can be influenced by two key prey functional traits: motility and defensive clumping. Strain *vfl1-1* has impaired swimming ability compared to WT (Adams *et al*. 1985;Appendix V videos) and is predicted by theory to experience lower predation rates relative to the more motile WT, all else equal (Aljetlawi *et al*. 2004; González *et al*. 1993; Pawar *et al*. 2012; Visser 2007). Our data supports this prediction as *vfl1-1* is selected against in the absence of predators, and selected in favor in the presence of most predators (Fig. 2 c-d) owing to decreased predation through impaired motility relative to WT.

However, when predation pressure is high enough, WT can develop plastic defensive clumps (Lurling & Beekman 2006), while *vfl1-1* cannot. We quantified the onset of this clumping defensive trait as cytometric changes in algal particle size over time: a faster increase in average particle size over time should indicate a quicker onset of defensive clumping. Genotype *vfl1-1* showed no significant clumping in response to predation (Fig. 2e yellow, Appendix IV Table 3) while WT showed strong plastic clumping against predation (Fig. 2e green; Appendix II Fig. 3-4; Appendix IV Table 4). The magnitude of the clumping, as measured by the shape of the distribution of WT particle size, varied by predator (Fig. 2e, Appendix IV Table 4), indicating predator-specific clumping responses likely resulting from model-predicted differences in predation rates. Last, we found differences in the relative fitness of motility and defensive clumping traits that are predator-specific. For example, *Glaucoma sp.* selected strongly in favor of *vfl1-1* while predator *P. caudatum* drove *vfl1-1* to extinction (Fig. 2c-d). These results show that food web context not only influences prey evolution, but also determines how prey traits shape the evolutionary process itself.

### Temperature only affects eco-evolutionary outcomes in the presence of predation

Temperature alone increased both WT and *vfl1-1* growth rates (Appendix II Fig. 5, Appendix IV Table 5), but did not affect prey rapid evolution –measured as rapid shifts in genetic frequencies– on its own (Fig. 3a Control panel; Appendix II Figure 2). However, temperature significantly influenced prey rapid evolution in the presence of predation, indicating that the temperature effects on prey evolution are mediated by predation (Fig. 3a; Appendix IV Table 6). We propose two concurrent mechanisms to explain this result: first, temperature influences prey evolution through differences in predator thermal performance and predation rates, leading to changes in selection on prey phenotypes (Fig. 3d mechanism 1). Second, temperature can affect prey evolution through its effect on the timing and magnitude of prey plastic responses induced by predation (Fig. 3d mechanism 2). Importantly, in the absence of predators, neither mechanism is at play, explaining why temperature does not drive prey evolution in this system on its own, and how food web context can influence the evolutionary outcome.

**Figure 3.**
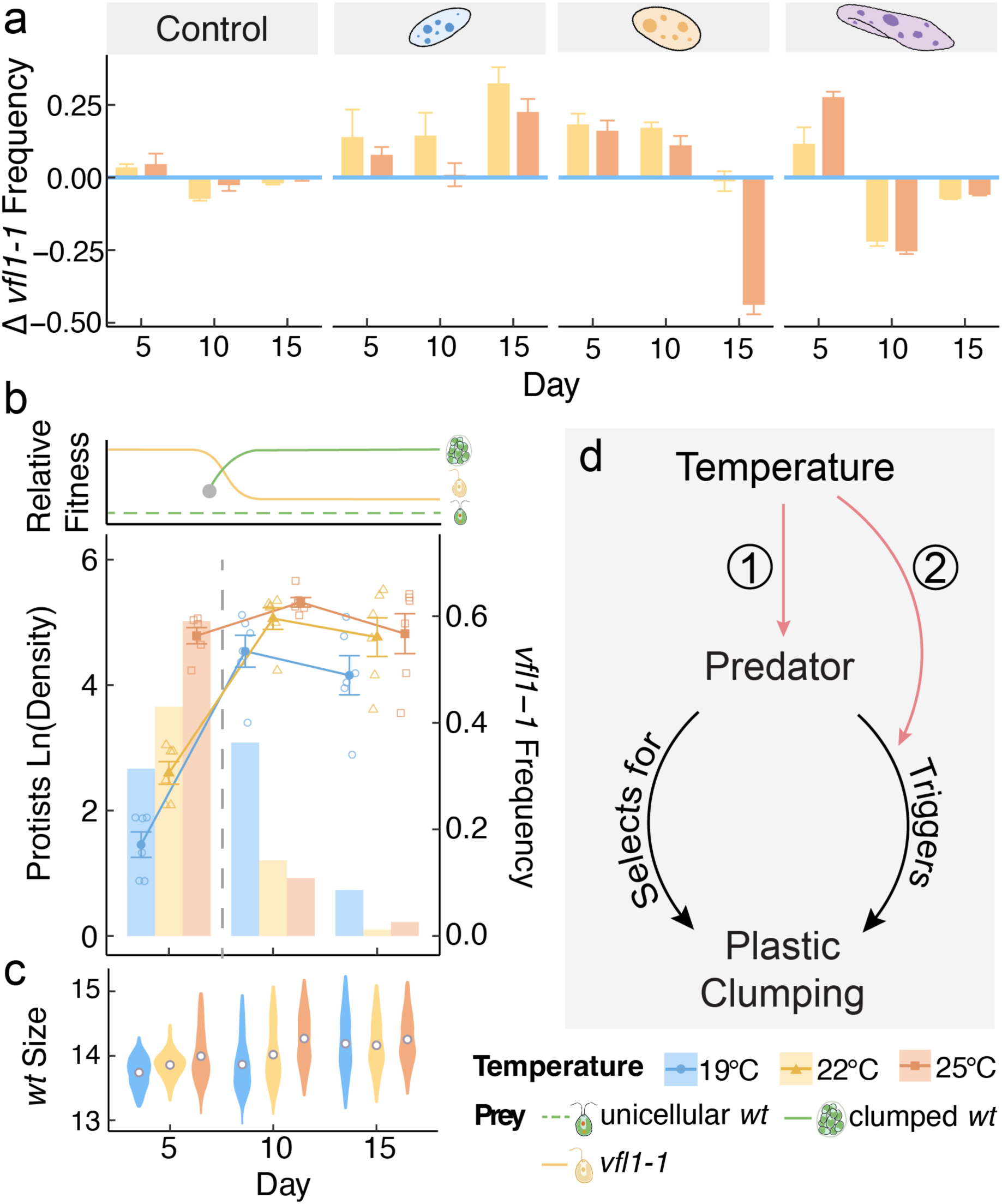
Temperature affects prey evolution through predator thermal performance and prey plasticity. a) To understand temperature effects on prey evolution, we calculated the changes in *vfl1-1* frequencies across temperature in control and all single predator treatments, using *vfl1-1* frequency at 19°C as the baseline (representing as blue horizontal line at 0). Blue, yellow, and orange represent 19°C, 22°C, and 25°C respectively. b) shows the *P. caudatum* density (left axis, solid lines) and *vfl1-1* frequency (right axis, bars) across temperature in *P. caudatum* single predator treatment. Circles, triangles, and rectangles represent 19°C, 22°C, and 25°C, respectively. The grey dash line represents the switch in the selection direction after plastic response of WT. Top panel is a conceptual diagram indicating the decrease of relative fitness of *vfl1-1* (yellow solid line) due to the onset of WT clumping (grey dot). The relative fitness of unicellular WT and clumped WT are presented by green dash and green solid line, respectively. c) WT particle size distribution in *P. caudatum* treatment across temperature. d) Conceptual diagram of the two potential mechanisms of temperature affecting WT clumping.

To illustrate how these mechanisms explain the observed eco-evolutionary dynamics, we focus on predation by *P. caudatum* as a case study. The first mechanism was evidenced by temperature-dependent changes in the predator demographic parameters that underpin their population dynamics: the intrinsic growth rate, *r* (Appendix II Fig. 6, Appendix IV Table 7), and predator maximum abundances, which serve as a proxy for the carrying capacity, K (Fig. 3b, Appendix II Fig. 6, Appendix IV Table 7). Both r and K are tied to predation intensity, as faster population growth (r) and larger carrying capacity (K) require increased predation to fuel and maintain biomass growth. Consequently, increased predation accelerated prey evolutionary dynamics by exerting stronger selection on the prey, resulting in increased predation rates and stronger selection against WT for its high motility in the early stage of the experiment (or, conversely, in favor of *vfl1-1*, Fig. 3b data panel, bars left of dash line; Appendix IV Table 2, 6). Evidence for the second mechanism comes from the differences in the timing and magnitude of WT plastic responses across temperatures (Fig. 3c and Appendix II Fig. 3). With the onset of plastic clumping (Fig. 3b, grey dot in conceptual panel), WT had higher relative fitness than *vfl1-1* in the later stages (Fig. 3b conceptual panel), leading to a decline in the less-defended *vfl1-1* (Fig. 3b data panel, bars right of the dash line). Importantly, temperature-dependent predation rates (Fig. 3b) influence the clumping response through changes in the distribution of WT particle size, including mean, median, variance, skewness, and kurtosis (Fig. 3c; Appendix IV Table 8). The intensified WT plastic responses at higher temperatures (Fig. 3c) and the subsequent temperature-dependent drop in *vfl1-1* frequencies (Fig. 3b) further evidenced the effects of temperature on the anti-predator defense and ensuing eco-evolutionary dynamics (Fig. 3b).

### Predator competition modulates how predation affects prey evolution across temperatures

To understand the importance of the broader food web context, we introduced a second predator (i.e., a predator competitor) and assessed their effect on the observed eco-evolutionary dynamics across temperatures (Fig. 4). We did so by quantifying the similarity between prey evolutionary dynamics in single-predator vs predator-competition treatments using Earth Mover’s Distance (see Methods). First, we discovered that the directional selection applied by each predator individually is attenuated when two competing predators favor distinct strains (Fig. 4). Specifically, prey evolutionary dynamics of single predator treatments were least similar to one another between each pair of the competing predators (e.g. prey evolutionary dynamics of *Glaucoma sp.* and *P. caudatum* single predator treatments have high EMD score, or low similarity; Appendix III Fig. 1). In contrast, the evolutionary dynamics of competition treatments (e.g. *Glaucoma sp.* + *P. caudatum*) showed higher similarity to the evolutionary dynamics of either single predator treatments (Appendix III Fig. 1).

**Figure 4.**
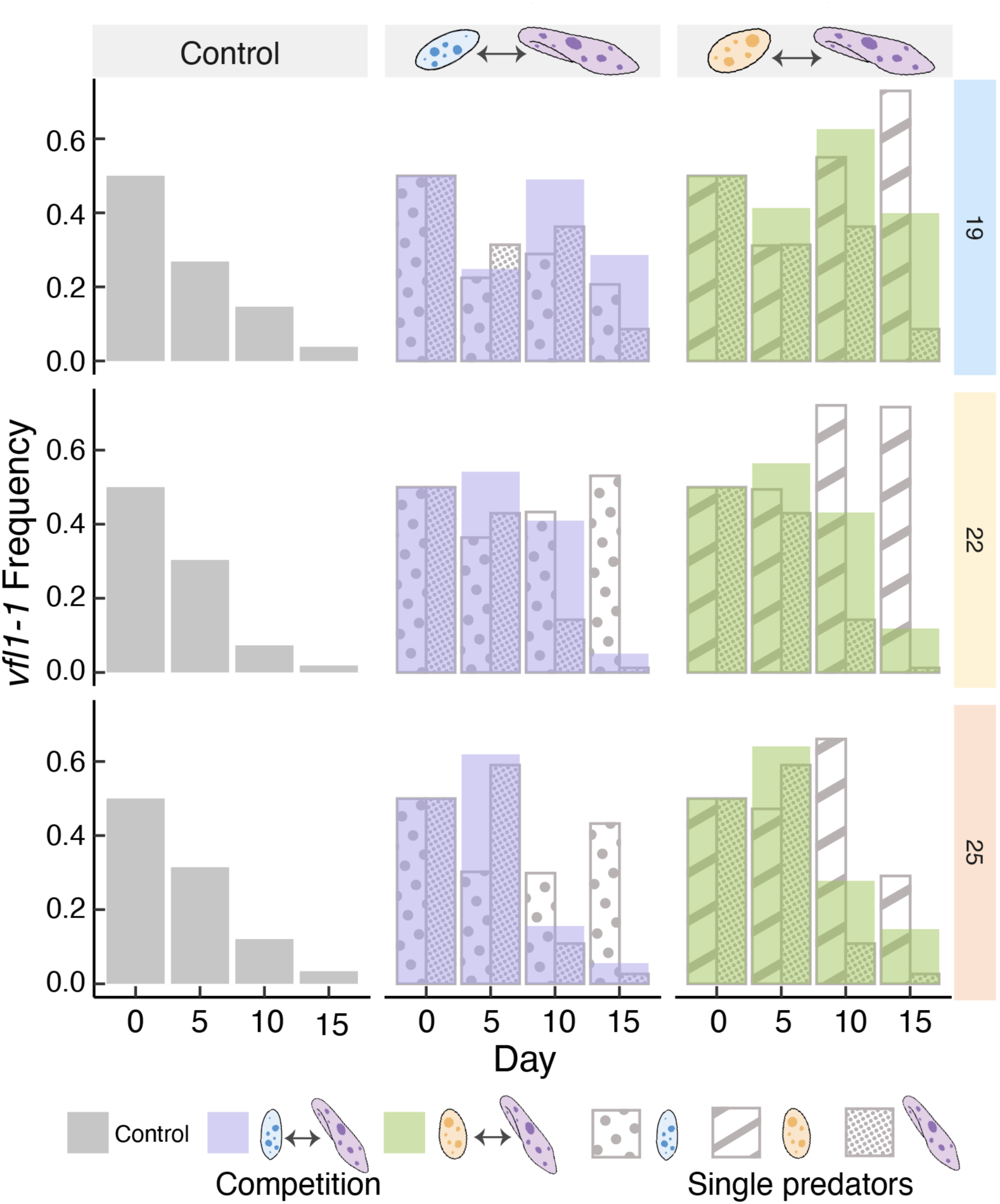
Prey evolution under predator competition. Frequencies of *vfl1-1* in competition treatments with two predators. Solid color bars represent the frequencies of *vfl1-1* in control and two predator-competition treatments at different temperatures. Bars with patterns represent the frequencies of *vfl1-1* in the single-predator treatments at each temperature.

Second, we found that predator competition mediated temperature effects on prey evolution (Fig. 4; Appendix IV Table 9-12) through the same mechanisms described previously (Fig. 3d). Particularly, differences in predator thermal performance drive differences in their relative contribution to selection on prey traits across competition treatments (Appendix II Fig. 7a), leading to temperature effects on prey evolution in the presence of predation and predator competition. For example, predation favored *vfl1-1* in all competition treatments where *T. pyriformis* and *Glaucoma sp.* had their highest thermal performance in competition with *P.caudatum,* (i.e. highest r and max population density at 19°C, Appendix II Fig. 6; Fig. 3d mechanism 1; Fig. 4) and WT showed the slowest clumping response (Appendix II Fig. 7b), resulting in strong selection against WT (Fig. 4; Appendix II Fig. 7a; Fig. 3d mechanism 2). However, at warmer temperatures, *P. caudatum* became more abundant and performed better (Appendix II Fig. 6), exerting stronger selection on prey than its competitors (Fig. 3d mechanism 1; Fig. 4). This resulted in faster and stronger plastic response by WT (Fig. 3d mechanism 2), selection in favor of WT, and accelerated loss of *vfl1-1* in competition treatments at high temperature (Fig. 4, 25°C; Appendix Fig. 7a). Consequently, prey evolutionary dynamics in competition treatments more closely resembled those in *T. pyriformis* and *Glaucoma sp.* single predator treatments at cooler temperatures, but more resembled that in *P. caudatum* treatment at higher temperatures (Appendix III Fig. 1). Taken together, these results demonstrate food web context influences the effects of temperature on these eco-evolutionary dynamics in a predictable way through the interplay between prey plasticity, predator thermal performance, and competitive ability within the broader food web. The density dynamics of both prey genotypes and all predator species across all treatments can be found in Appendix II Fig. 8.

## DISCUSSION

Our results reveal that temperature strongly but indirectly affects prey evolution within food webs through differences in temperature-dependent predation rates across genotypes and prey defensive plastic responses (Figs. 2-3). In the presence of multiple predators, differences in predator performance across temperatures leads to predator-specific differential selection across prey genotypes–which would not occur in the absence of predation and ultimately lead to differences in prey evolutionary outcomes across food web contexts (Fig. 4). We highlight the significance of the broader food web context in shaping species evolution under global warming, and show how even identical species in different food webs may respond to warming in different ways.

### Plasticity influences prey rapid evolution

Our study shows that predator differential selection on multiple prey functional traits (i.e. motility and clumping) facilitates the maintenance of additive genetic diversity (Fig. 2, Siepielski *et al*. 2020)). Moreover, we revealed the reciprocal effects of prey functional traits and species interactions, and how these underpin the dynamics of species evolution within food webs (Fig. 3). While previous studies have shown that more motile prey experience higher predation rates compared to slower prey (Andersen & Dölger 2019; González *et al*. 1993), we found that the relative fitness of the less motile *vfl1-1* strain was influenced by the onset of a plastic defensive trait in the WT strain, itself temperature-dependent (Fig. 3). The onset of this trait-mediated eco-evolutionary process sometimes led to a new equilibrium in which all genetic variants persisted, or one in which only one did (Fig. 3). Consequently, dynamic, temperature-dependent plastic responses can significantly alter the course of the evolutionary process and influence the fate of genetic variants in a warming world. However, this process is vastly underappreciated and poorly understood in food web ecology.

### Predator identity and food web context affect prey plastic responses

We showed that evolutionary outcomes are strongly dependent on predator identity and the broader food web context, as the interplay between rapid plasticity and evolution can shift predator preferences (Fig. 3). Additionally, multiple predators can have joint effects in shaping prey plastic and evolutionary responses (Fig. 4; Appendix III Figs. 1-2), which is itself contingent on the composition of the food web. Indeed, prey evolutionary dynamics under selection by multiple predators reflected the combined effects of selection imposed by individual predator species (Fig. 4; Appendix III Fig. 1). Phenotypic change, plastic or evolutionary, can thus determine the fate of organisms within food webs: shifts in one functional trait in response to one predator may affect the interaction with another predator, potentially cascading through the entire community, as had been predicted by theory (Cosmo *et al*. 2023; Guimarães *et al*. 2017). Our study highlights the importance of achieving a mechanistic understanding of the interplay between species functional trait dynamics and biotic interactions across trophic levels to better understand their fate in a changing world (Henn *et al*. 2018).

### Food web context and temperature interactively drive prey evolution

While temperature-mediated changes in species interactions have been hypothesized to drive changes in species plastic and evolutionary functional traits in a warming world (Barbour & Gibert 2021; Fischer *et al*. 2016; Fordyce 2006), empirical support for this hypothesis is lacking. We show that reciprocal effects between food web context and trait dynamics together drive prey evolutionary trajectories differentially across temperatures (Figs. 3-4). If food-web context and temperature jointly influence rapid changes in the genetic makeup of populations, as shown here, warming could severely impact species persistence across in warming food webs. For example, we show that predation and predator diversity can maintain prey trait diversity (Fig. 2). However, observed increasing extinction rates among predators under warming (Thunell *et al*. 2021; Zarnetske *et al*. 2012) may prompt a decrease in predator diversity and subsequent reduction in prey genetic and trait diversity, in turn decreasing potential for future adaptive evolution. This loss of genetic diversity could be most pronounced when dominant predators disproportionally affect prey evolution under warming (e.g., *P. caudatum*, Fig. 4), particularly in simpler food webs at higher latitudes (Gibert 2019). Standing plasticity and the evolution of plasticity have the potential to alter these outcomes, underlining the importance of understanding their ecological consequences in a rapidly changing world. Last, while species evolution is shaped by the food web context (Figs. 2-4), the composition and network complexity of food webs are rapidly changing, which should have important but currently unknown effects on evolutionary outcomes (Barbour & Gibert 2021).

## CONCLUSIONS

Our research reveals that temperature influences prey evolution exclusively in the presence of predators, highlighting the crucial role of food-web context in determining eco-evolutionary outcomes. While rapid evolution has been proposed as a potential buffer against the impacts of climate change on natural communities, our study underscores that anticipating evolutionary outcomes and species fate in novel climates demand a deeper understanding of evolutionary responses across food-web contexts and accounting for rapid shifts in these food webs as the globe warms.

## METHODS

### Experimental work

Unicellular microalgae can be found in diverse habitats, from soil to freshwater ecosystems worldwide (Arora & Sahoo 2015; Falkowski 1994). These organisms are at the base of all food webs, fueling both green and brown energy pathways (Guo *et al*. 2016), and are routinely preyed upon by microbes (e.g., ciliate protists) and metazoans (e.g., rotifers, cladocerans) alike (Calatrava *et al*. 2023). Our focal prey species, the unicellular green alga *Chlamydomonas reinhardtii*, is a well-established model organism with known mutations linked to functional traits (Calatrava *et al*. 2023; Sasso *et al*. 2018; Tulin *et al*. 2024). Here we used two genetically and phenotypically distinct strains of *C. reinhardtii*: wild type (WT) and *vfl1-1* (*v*ariable *fl*agella) as our prey population.

Wild type (WT) possesses two flagella that allow the entire range of normal locomotive behaviors (Huang 1986) and can form large cell clumps, a common form of defense against predation (Herron *et al*. 2019). Strain *vfl1-1* produces individuals with variable numbers of flagella from 0 to 10 and defective swimming ability (Adams *et al*. 1985). Additionally, *vfl1-1* seems unable to clump under predation. To distinguish these two strains and keep track of their relative frequencies in mixture populations, we used mNeonGreen fluorescent protein to tag WT populations and used flow cytometry to count tagged WT vs non-tagged *vfl1-1* individuals over time (Fig. 1a). Both strains exhibit autofluorescence (Fig. 1a), but only one (WT) will show fluorescence in the near green spectrum (Fig. 1a). We used two protist predators of similar body size, *Tetrahymena pyriformis* and *Glaucoma sp.*, as the focal predators, and a larger protist, *Paramecium caudatum*, as a competitor for the focal predators (Fig. 1b). All four species are commonly found in freshwater and soil systems (Cornwallis *et al*. 2023; Foissner & Berger 1996).

To understand how temperature and food web context (i.e., presence of predators and predator competitors) jointly influence *C. reinhardtii* evolution, we set up experimental microcosms in autoclaved 250 ml borosilicate jars filled with 100 ml of 9:1 COMBO media (Kilham *et al*. 1998): timothy hay infusion (Brans *et al*. 2022). Each microcosm was assigned to one of three possible predation treatments (no predation/Control, + *T. pyriformis*, + *Glaucoma sp.*), one of two possible predator competition treatments (+ *T. pyriformis* & *P. caudatum*, + *Glaucoma sp.* & *P. caudatum*), and one of three possible temperatures: 19°C, 22°C, and 25°C. The manipulations produced a factorial design with 18 combinations of treatments (Fig. 1b), each replicated six times, yielding a total of 108 microcosms.

Prior to experimentation, the algae strains were maintained on TAP agar (Gorman & Levine 1965) at room temperature. Protist cultures were maintained in bacterized timothy hay protist media at 22°C and a 16:8-hour light-dark cycle. All cultures were transferred to 9:1 COMBO media: timothy hay protists media and cultured under the same light and temperature regime as protist stock cultures 2 weeks prior to experimental work. We carried out the experiment in two blocks on two consecutive days; each block had half of the replicates in all treatments. We started WT and *vfl1-1* strains at equal densities of 2000 individuals/ml in all microcosms and initialized *T. pyriformis*, *Glaucoma sp.*, and *P. caudatum* populations at density of 5 ind/ml, 5 ind/ml, and 0.5 ind/ml, respectively. The experiment was carried out for 15 days, or ∼ 30-45 *C. reinhardtii* generations under control conditions at the three focal temperatures.

We recorded all species densities on days 0, 5, 10, and 15. We used flow cytometry (NovoCyte 2000R, Agilent, CA, USA) to distinguish the two genetically and phenotypically distinct strains of *C. reinhardtii*: mNeonGreen tagged wild type (WT) and *vfl1-1* (*v*ariable *fl*agella). This allowed us to track the abundances and frequencies of both strains over time. We used forward scatter height (FSC-H) as a measure of cell/cell clump size of *C. reinhardtii* (Adan *et al*. 2017) to track plastic morphological change in response to protist predation, thus providing a window into both rapid plastic change (i.e., within strains), and rapid evolutionary change by clonal sorting (i.e., change in genetic frequencies; Fig. 1c). We recorded the density of all protist species through fluid imaging (FlowCam; Yokogawa Fluid Imaging Technologies, Portland, ME, USA) at a magnification of 10x (Fig. 1c).

### Data analysis

To analyze how temperature and ecological interactions influenced prey evolution, we used autoregressive moving average linear mixed models (ARMA-LMMs; ‘nlme’ package, v. 3.1-162) to examine autocorrelation in our data. After finding no significant effects of autocorrelation (Appendix IV Table 1), we used classic linear mixed models (LMM) in the ‘lme4’package (version 1.1-3 in R v. 4.3.1) to the rest of analyses. We calculated the relative frequencies of *vfl1-1* in total *C. reinhardtii* population as the measure of prey evolution. To better evaluate whether changes in prey evolution were affected by temperature, we also calculated the relative changes in *vfl1-1* frequencies across temperatures (Δ *vfl1-1* frequency) by deducting the mean *vfl1-1* frequencies at 19°C from each replicate at 22°C and 25°C in each treatment.

We first performed LMM on the single predator treatments and the control (Fig. 1b) to understand the individual effects of each predator on prey evolution. We analyzed the fixed effects of predator species identity, temperature, and time on the prey evolution and temperature effects on prey evolution, by using *vfl1-1* frequency and Δ *vfl1-1* frequency as response variables respectively. We then accounted for the densities of the single predators as a fixed effect to the previous model to analyze the effects of predation on prey evolution. We then performed LMM on *T. pyriformis* and *Glaucoma sp.* single predator treatments and the competition treatments (Fig. 1b) to test whether the presence of a second predator had joint effects with predator species identity, predator density, and temperature on *vfl1-1* genetic frequency and Δ *vfl1-1* frequency using previous linear mixed models.

Additionally, to quantify the relative importance of each predator on prey evolution in competition treatments, we calculated Earth Mover’s Distance (EMD from now on) with ‘emdist’ (R package, version 0.3-3), which quantifies the similarity of prey evolution patterns over time between different predation treatments (higher EMD means less similarity). Specifically, at each temperature, we calculated the EMD of *vfl1-1* frequencies 1) between the single predator treatments in each competition pair and 2) between each of the single predator treatments and the corresponding competition treatment.

To understand the mechanism through which predation and temperature jointly affect prey evolution, we quantified demographic parameters that govern species population dynamics: 1) a proxy for maximum growth rate, r, at each temperature, as ln(N_t_)−ln(N_0_)]/time using day 5 data and their initial densities on day 0, and, 2) maximum density, N_max_, by measuring the highest daily average across replicates. We used linear models (‘stats’ v4.3.1) and stepwise model selection (‘stepAIC’ in R package ‘MASS’ v7.3-60) to test the effects of temperature and competition on each of the predators.

We also used linear mixed models to understand how abiotic and biotic factors affect prey plasticity. Similar to previous LMMs, we tested how temperature, predator species, predator density, and time, affect WT and vfl1-1 particle sizes in single predator treatments. In addition to those factors, we also tested how the presence/absence of a predator competitor affects prey particle size in the competition treatment.

### Mathematical modeling

To understand the processes that drive rapid evolution in the prey population in the presence/absence of a predator, we mathematically tracked the population dynamics of a system with two genetically distinct prey strains under predation by a shared predator. We assumed that WT (W) and *vfl1-1* (V) grow logistically and compete for resources at different rates using a model in differential equations. Further, we assumed that the predator (P) has a multispecies type II functional response and dies at a background mortality rate, *m*. Our model allows for the predator to prey on each strain at different rates. Taken together, the equations modeling the strain dynamics and predator population are:

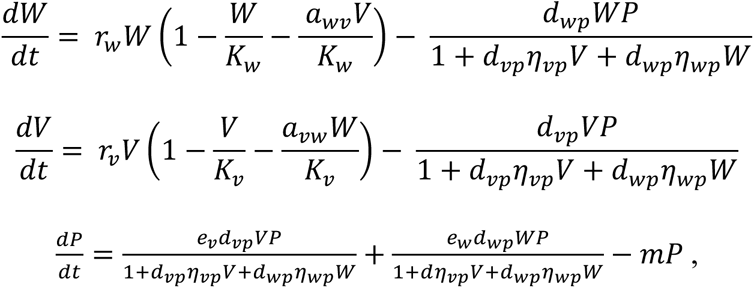

where r is intrinsic growth rate, K_i_ is the carrying capacity of each genotype, *a_wv_*is the competition effect of V on W (vice versa for *a_vw_*), *d_wp_* and *η_wp_* are the attack rate and the handling time of the predator on W, *a_vp_* and *η_vp_* are those of predator on V, *e_w_* and *e_v_* are the conversion efficiencies of W and V to predator biomass. We used the growth rate of WT and *vfl1-1* in control conditions as *r_w_* and *r_v_* and then explored parameter space with the remaining model parameters to find dynamics that qualitatively reproduced the observed dynamics. In the appendix we also include alternative model formulations (e.g., treating predation as a constant mortality rate, predators with a type I functional response) and provide an analytical treatment of the model and associated predictions (Appendix I).

