## Supplemental Materials for "Temperature alone is not enough: food-web context determines evolutionary responses to warming"

**Extended methods**

**Appendix I** Additional Mathematical Models …………………………………………………...2

**Appendix II** Species body size, ecological, and evolutionary dynamics…………..…………...6

**Appendix III** Earth Mover's Distance……….……………………………………………….12

**Appendix IV** Statistical analysis…………….……………………………………………….14

### Appendix I Mathematical Models

For all models, we assume *Chlamydomonas reinhardtii* wild type (WT) and mutant strains *vfl1-1* grow logistically with different growth rates, $r_w$ and $r_v$, respectively. These two strains are assumed to compete for resources with competition coefficients $a_{wv}$ and $a_{vw}$, where $a_{wv}$ represents the effects population WT has on population *vfl*, and vice versa for $a_{wv}$.

#### *1. Predation as a constant rate*

We first assume predation affects the growth of each strain at a constant rate on a per capita basis (i.e., a constant proportion of the population is being removed by the predator per unit time). We also assume that the predators remove WT and *vfl1-1* at different rates, $p_w$ and $p_v$. Therefore, the changes in the density of WT (W) and *vfl1-1* strain (V) strains can be written as:

$$\frac{dW}{dt} = r_w W\left(1 - \frac{W}{K_w} - \frac{a_{wv}V}{K_w} - p_w\right)$$

$$\frac{dV}{dt} = r_v V\left(1 - \frac{V}{K_v} - \frac{a_{vw}W}{K_v} - p_v\right)$$

where $K_w$ and $K_v$ are the carrying capacity of WT and *vfl1-1*, respectively, and $p_w$andf $p_v$ are the constant predation rate on WT and *vfl1-1.*

For *vfl1-1* and WT to coexist, both *vfl1-1* and WT need to meet the invasibility criterion, such that each has to be able to invade the system when in low abundance and while the other species is at its equilibrium (i.e. carrying capacity if $p_i$=0). For *vfl1-1* to meet the invasibility criterion, $\lim_{V\to 0} \frac{dV}{dt} > 0$ at the equilibrium of (W*, 0), i.e., with $W* = K_w(1 - p_w)$. Under these conditions, *vfl1-1* invades when:

$$\frac{K_v}{K_w} > \frac{a_{vw}\,(1-p_w)}{1-p_v}. \quad [1]$$

For WT to invade the system when rare, the condition $\frac{K_w}{K_v} > \frac{a_{wv}\,(1-p_v)}{1-p_w}$ has to be met. We notice that when predation is weak ($p_i$~0), the conditions become

$$\frac{K_v}{K_w} > a_{vw}$$

$$\frac{K_w}{K_{vw}} > a_{wv} \text{ , [2]}$$

which are the classical conditions for coexistence under competition. However, with predation, *vfl1-1* can invade whenever one or more of the following criteria are met: 1) carrying capacity of V is large, the carrying capacity of W is small, or both; 2) the competition coefficient of W on V, $a_{vw}$, is small; 3) predation on W, $p_w$, is large, while 4) predation on V, $p_v$, is small.

***2. Predator with type I function response***

The previous model did not include density dependence in the predation process. In this second model iteration we explicitly do by accounting for changes in predator density (P). We assume the predator preys on WT (W) and *vfl1-1* (V) at different rates with a type I functional response and has a constant mortality rate, $m$. The dynamics of 2-prey-strain-and-predator system can be written as:

$$\frac{dW}{dt} = r_w W\left(1 - \frac{W}{K_w} - \frac{a_{wv}V}{K_w}\right) - d_{wp}WP$$

$$\frac{dV}{dt} = r_v V\left(1 - \frac{V}{K_v} - \frac{a_{vw}W}{K_v}\right) - d_{vp}VP$$

$$\frac{dP}{dt} = e_w d_{wp}WP + e_v d_{vp}VP - mP$$

where new parameters $d_{wp}$ and $d_{vp}$ represent the attack rate of the predator on WT and *vfl1-1* respectively, $e_w$ and $e_v$ represents the conversion efficiency of the prey strains WT and *vfl1-1*, respectively, into predator biomass.

In this model, coexistence between WT and *vfl1-1* without the predator is possible (W* = $\frac{K_w - a_{wv}K_v}{a_{vw}a_{wv}-1}$, V* =$\frac{K_v - a_{vw}K_w}{a_{vw}a_{wv}-1}$, P* = 0). However, for *vfl1-1* to invade the system when rare and with V and P at equilibrium (i.e., W* = $\frac{m}{d_{wp}e_w}$, V* =0, P* = $\frac{(d_{wp}e_w K_v - m)r_v}{{d_{wp}}^2\, e_w\, K_w}$), the invasion criteria is

$$K_v > a_{vw}\frac{m}{d_{wp}e_w}. \text{ [3]}$$

This indicates that for *vfl1-1* to invade the WT and predator two-species equilibrium, the carrying capacity of *vfl1-1* ($K_v$) needs to be larger than the product of inter-clonal competition between prey strains and the ratio between predator mortality and predator conversion efficiency times its attack rate on WT. For this to be true, one or more of the following should happen: 1) the inter-clonal competition, $a_{vw}$, is low (i.e., WT is a weak competitor); 2) predator mortality is low (likely keeping WT at bay); 3) predator attack rate on WT, $d_{wp}$, is high; 4) the conversion efficiency, $e_w$, is high, or a combination of all of the above. In other words, the invading strain needs to experience low competition from the resident strain and the resident strain needs to experience relatively high predation rates by an efficient predator, hence supporting the notion that predation maintains coexistence of the strains, and thus, genetic diversity. A similar argument is true for invasion of WT.

##### *3. Predator with multispecies type II function response*

We now add constrains on the predator intake rate by assuming the predator has a multispecies type II functional response. The dynamics of the system can be modeled as:

$$\frac{dW}{dt} = r_w W\left(1 - \frac{W}{K_w} - \frac{a_{wv}V}{K_w}\right) - \frac{d_{wp}WP}{1 + d_{vp}\eta_{vp}V + d_{wp}\eta_{wp}W}$$

$$\frac{dV}{dt} = r_v V\left(1 - \frac{V}{K_v} - \frac{a_{vw}W}{K_v}\right) - \frac{d_{vp}VP}{1 + d_{vp}\eta_{vp}V + d_{wp}\eta_{wp}W}$$

$$\frac{dP}{dt} = \frac{e_v dVP}{1 + d_{vp}\eta_{vp}V + d_{wp}\eta_{wp}W} + \frac{e_w d_{wp}WP}{1 + d\eta_{vp}V + d_{wp}\eta_{wp}W} - mP$$

where the new parameters $\eta_{wp}$ and $\eta_{vp}$ represent the handling time of the predator on WT and *vfl1-1* respectively. This is the version of the model used to reproduce the observed eco-evolutionary dynamics in main text Figure 2a-b.

We hypothesize that the key parameters that determine differential selection patterns between the three predators (i.e. *Tetrahymena pyriformis*, *Glaucoma sp.*, and *Paramecium caudatum*) are their attack rates on the strains WT and *vfl1-1.* Thus, we only vary the attack rate (bolded, Table1), $d_{wv}$ and $d_{vw}$, between each species to reproduce the observed dynamics. We parametrized $r_w$ and $r_v$ by calculating the average population growth rate of WT and *vfl1-1* in control treatment across all replicates regardless of temperature (italicized, Table 1). The parameters used to generate each qualitative model predictions are listed in Table 1.

*Table 1 Model parameters for main text Figure 2a-b.*

| Parameters | Control | Tetrahymena | Glaucoma | Paramecium |
|---|---|---|---|---|
| $W_0$ | 10 | 10 | 10 | 10 |
| $V_0$ | 10 | 10 | 10 | 10 |
| $P_0$ | NA | 2 | 2 | 2 |
| $r_w$ | *0.543* | *0.543* | *0.543* | *0.543* |
| $r_v$ | *0.363* | *0.363* | *0.363* | *0.363* |
| $a_{vw}$ | 1.3 | 1.3 | 1.3 | 1.3 |
| $a_{wv}$ | 0.35 | 0.35 | 0.35 | 0.35 |
| $d_{vp}$ | | **0.02** | **0.025** | **0.1** |
| $d_{wp}$ | | **0.08** | **0.12** | **0.12** |
| $K_w$ | 50 | 50 | 50 | 50 |
| $K_v$ | 50 | 50 | 50 | 50 |
| $\eta_{\mathrm{wp}}$ | | 0.08 | 0.08 | 0.08 |
| $\eta_{\mathrm{vp}}$ | | 0.08 | 0.08 | 0.08 |
| $e_w$ | | 0.1 | 0.1 | 0.1 |
| $e_v$ | | 0.1 | 0.1 | 0.1 |
| $m$ | | 0.1 | 0.1 | 0.1 |

### Appendix II Species ecological dynamics and body size dynamics

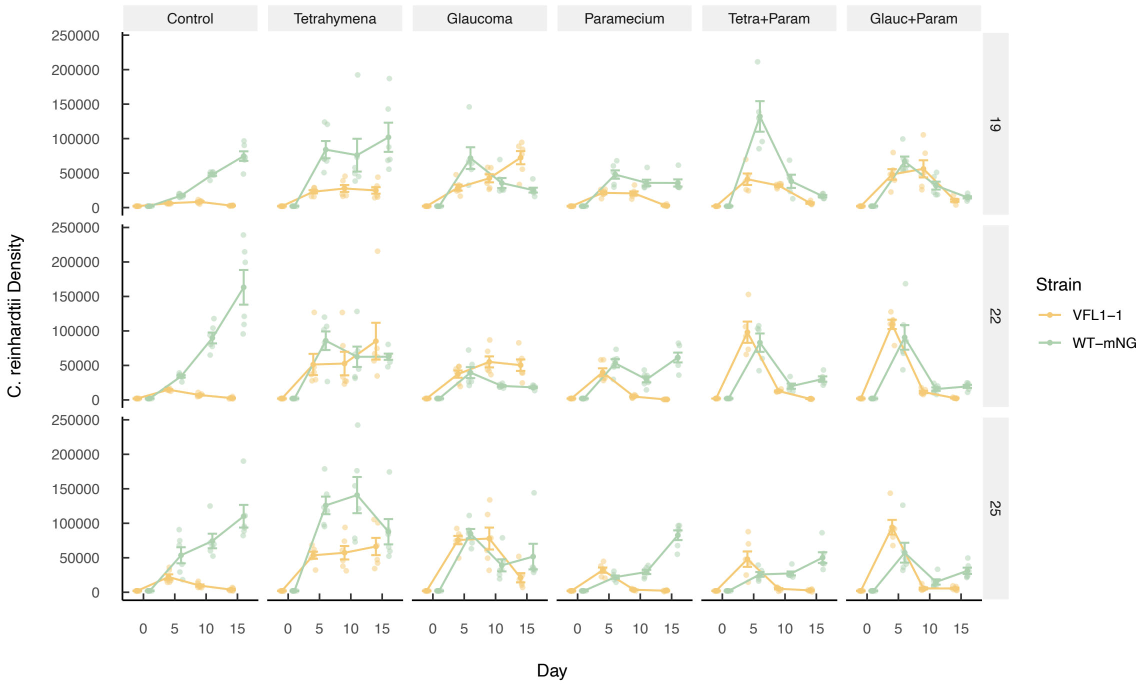


**Figure 1.** Population density of prey *Chlamydomonas reinhardtii* WT and *vfl1-1* across all predation and temperature treatments.

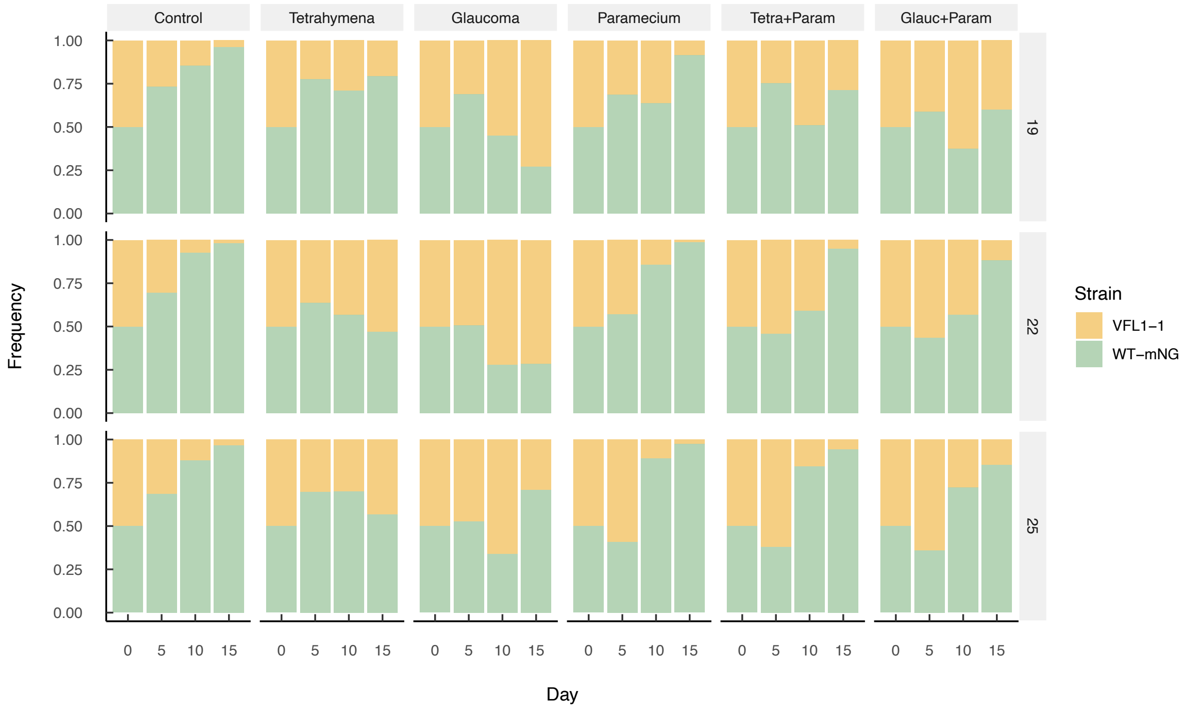


**Figure 2.** Clonal Frequency of vfl1-1 and wild type (WT). Column names represent the treatment name while the row name represent the temperature.

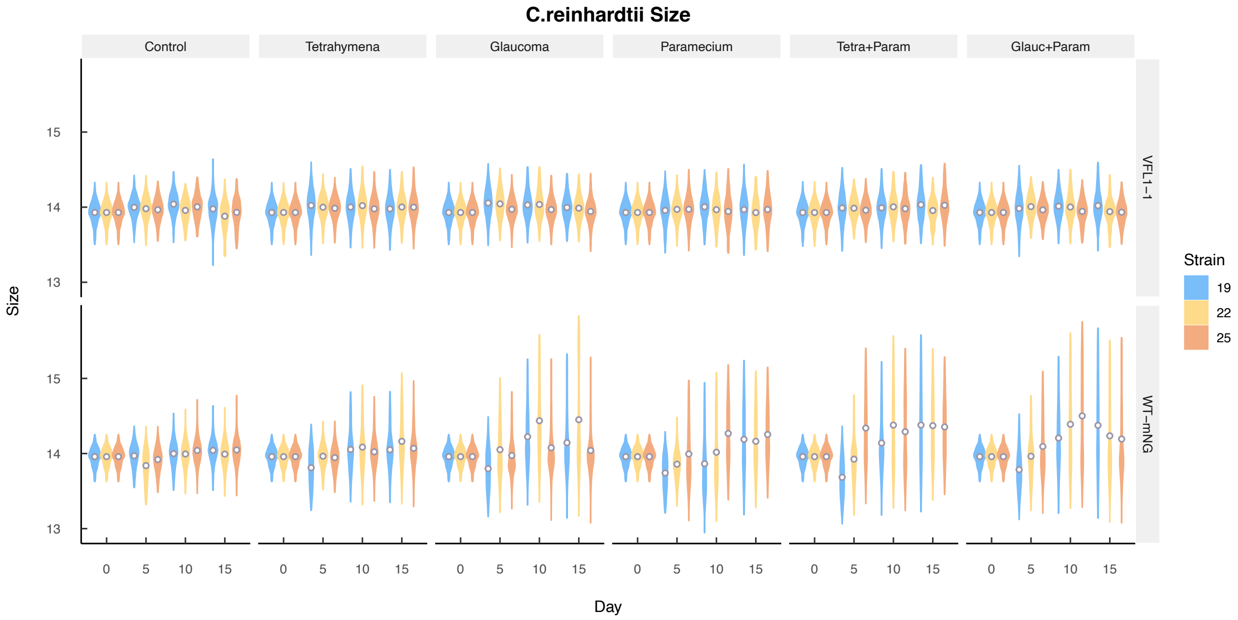


**Figure 3.** Cell clump sizes distribution of *Chlamydomonas reinhardtii* mutant strain *vfl1-1*(top row) and wild type (bottom row) across all treatments. Colored by temperature treatment.

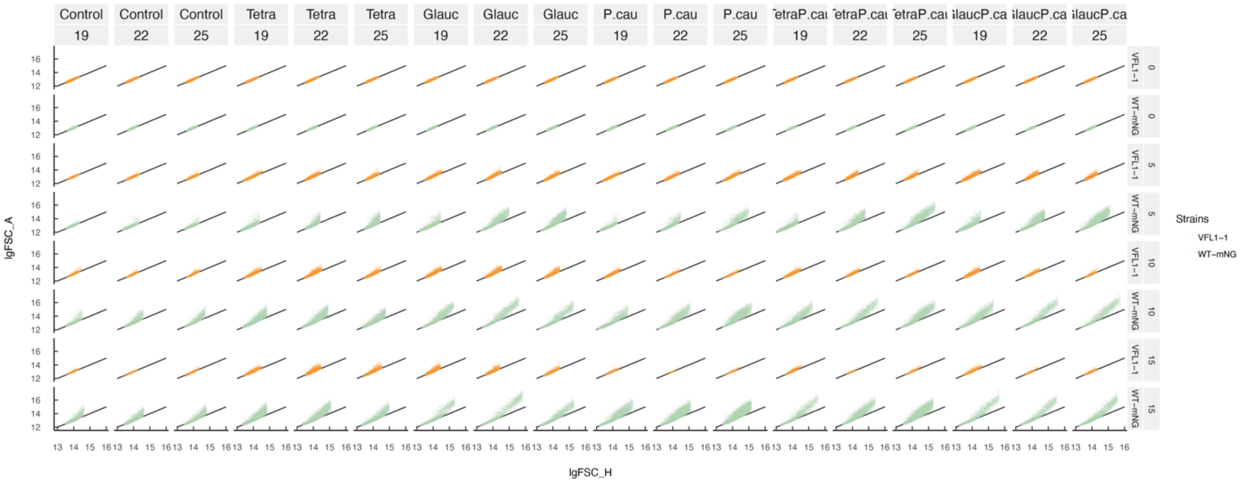


**Fig 4.** *Chlamydomonas reinhardtii* particle log forward scatter height (lgFSC_H) ploted against logged forward scatter area (lgFSC_A) to detect doublets (clumping). An increase in FSC_H does not necessarily mean multiple cellular clumpings. When the ratio of FSC_A:FSC_H of cells remains close to one, it suggests that only cell size are changing rather than the formation of multicellular clumps. When the ratio of FSC_A:FSC_H deviates from one, this indicates formation of multicellular clumps. We partition the cell size by all treatments, temperature, day, and strain to show the the ratio of FSC_A:FSC_H of each strain. WT is shown in green and *vfl1-1* in orange. Black line is a line with slope of one for reference. As we predicted, WT shows much larger range of FSC_H and its FSC_A:FSC_H ratio deviates significanlty from the 1:1 line, especially at larger FSC_H values. These results thus validate our hypothesis that WT formed more and larger multicellular clumps compared to *vfl1-1*.

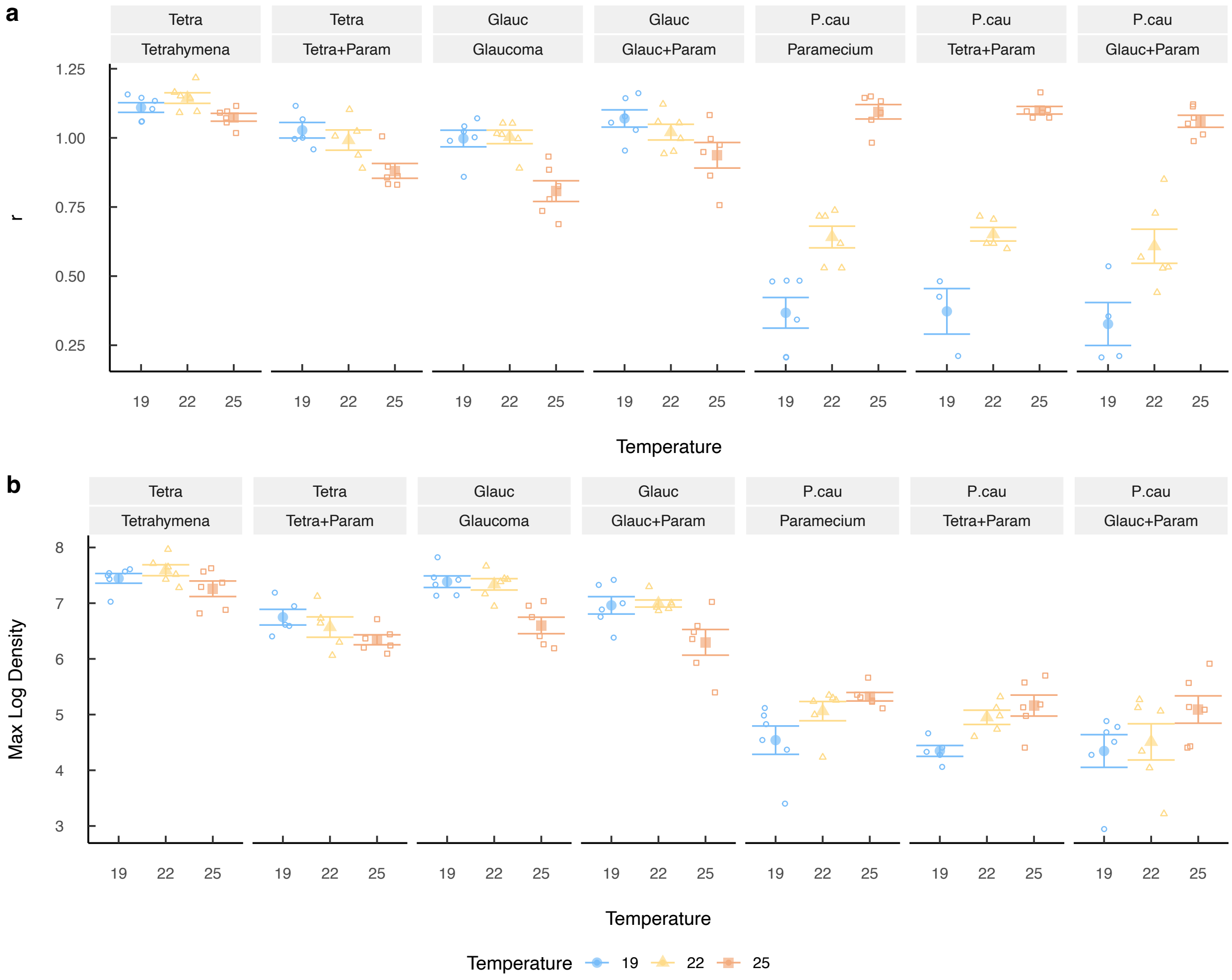


**Figure 5.** Initial population growth rate, r, at different temperature for WT and *vfl1-1* across all treatments. Initial population growth rate is calculated at each temperature, as $\ln(N_t) - \ln(N_0)$]/time using day 5 density data and initial densities on day 0. The average population growth rate of WT and *vfl1-1* across all replicates in control, regardless of temperature, is used for model parametrization in Appendix I Table 1. Blue circle represent 19°C, yellow triangles represent 22°C, and red rectangles represent 25°C. Solid shapes represent the mean, and open shapes represents individual replications.

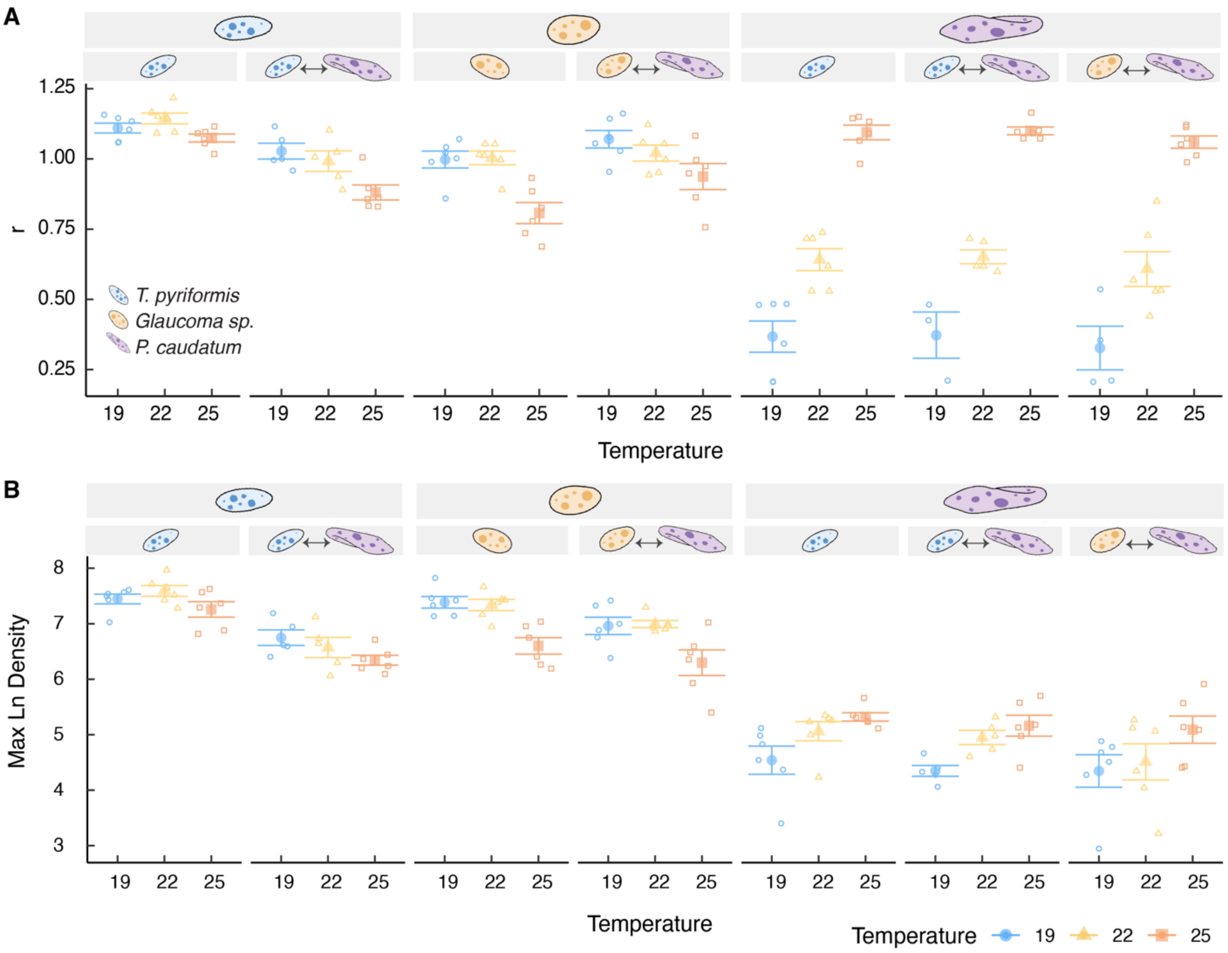


**Figure 6.** Initial population growth rate, r, and maximum abundance of each protist predator at different temperature across all treatments. Top panel labels represent species, and the bottom ones represents the treatments the protists were in. Methods and color scheme used are consistent with those in Fig 6.

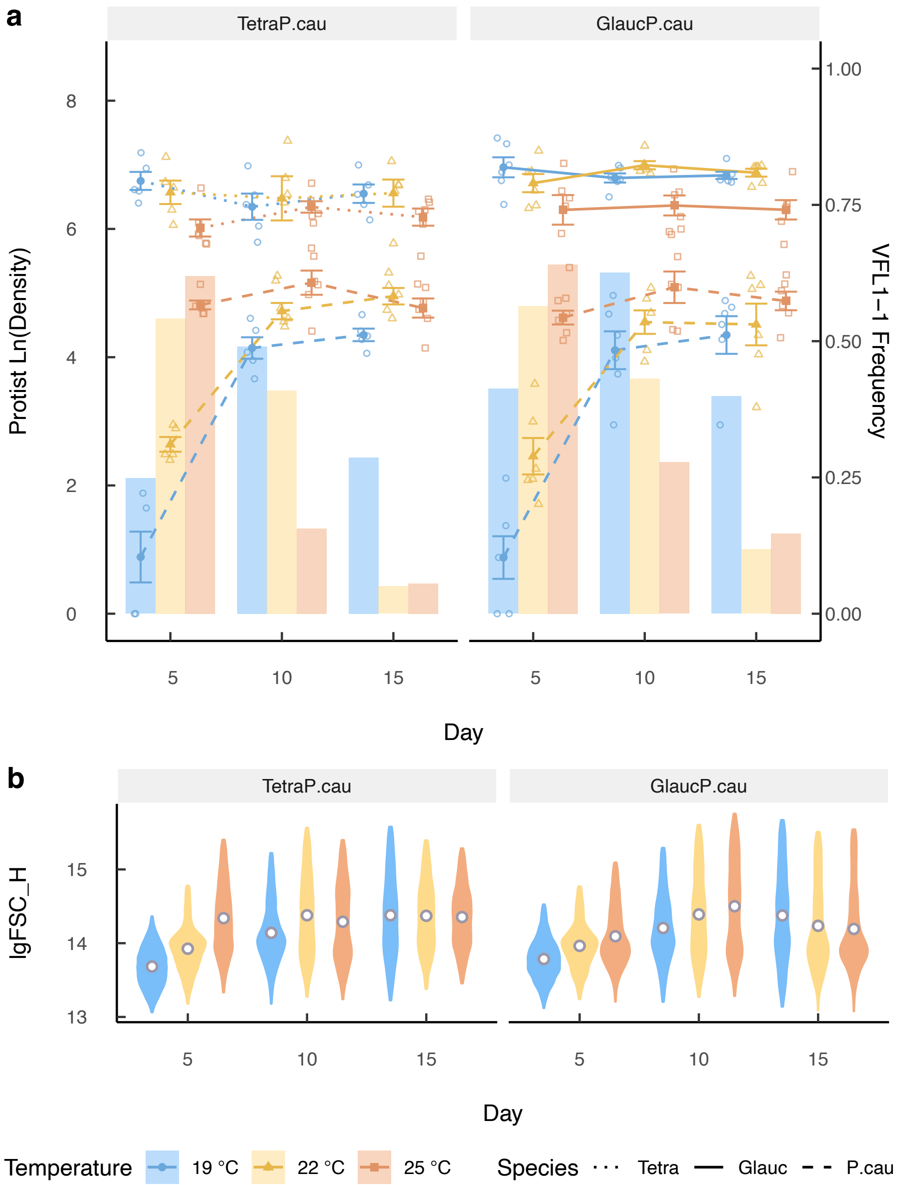


**Fig 7.** a) all predator log density (left axis, lines) and *vfl1-1* frequencies (right axis, bars) with error bars at different temperature. Triangles, squares, and circles represent 19°C, 22°C, and 25°C respectively. Dash line, dotted line, and solid line represent *T. pyriformis*, *Glaucoma sp.*, and *P. caudatum* respectively. Open shapes represent each replicate while the solid shapes represent the mean within each treatment. b) Wild type size distribution measured as log FSC-H at different temperature over time. White triangles and grey open circles represent the mean and the median.

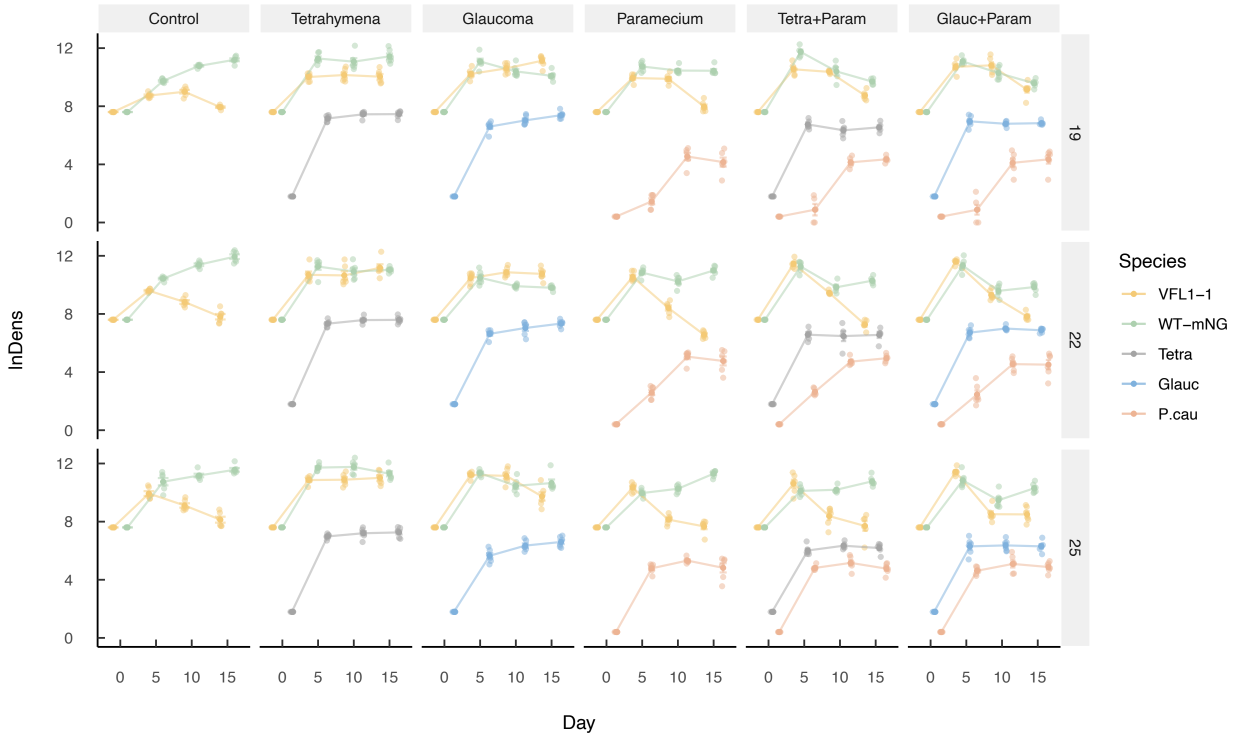


**Figure 8.** Log density of all species at different temperature treatments. Columns all the predator and competition treatments. Rows represent temperature treatments. Each color line represents the population density of *Chlamydomonas reinhardtii vfl1-1* mutant strain and WT as well as all the predators present in each treatment.

### Appendix III Earth Mover's Distance

In order to quantify the relative importance of each predator on prey evolution within predator competition treatments, we calculated Earth Mover's Distance (EMD from now on) with 'emdist' (R package, version 0.3-3), which quantifies the similarity of prey evolution patterns over time between different predation treatments (higher EMD means less similarity, Figure 1). Specifically, within each temperature, we calculated the EMD of *vfl1-1* frequencies 1) between the single predator treatments in each competition pair and 2) between each of the single predator treatments and the corresponding competition treatment. Similarly, to quantify the relative effects of each predator on prey plasticity, we also calculated the EMD of WT particle size distribution between: 1) the single predator treatments in each competition pair, and 2) each of the single predator treatments and the corresponding competition treatment (Fig 2).

In Figure 1, we found that, in general, the evolutionary dynamics of prey under the selection of single predator differ the most, as indicated with the highest EMD, and thus, least similarity (orange dotted line, Figure 1). However, under competition treatment where both predators are present, the prey evolutionary dynamics appear more similar to both single predator treatments (as shown by the grey and green lines in Figure 1). This suggests that in competition treatments, both predators play a role in determining the evolutionary outcome of the prey.

Interestingly, with increasing temperature, we observed an increasing similarity (decreasing EMD) in prey evolutionary outcomes between the Paramecium-only and both competition treatments, especially notable in the Glaucoma and Paramecium competition treatment (as represented by the grey solid line in Figure 1B). This suggests that when Paramecium has the highest temperature performance (i.e., highest growth rate $r$ and maximum population density $K$) while Tetrahymena and Glaucoma had the lowest performance at 25°C (Appendix II Fig 7), Paramecium has a stronger influence in driving prey evolution. However, at lower temperature, where Tetrahymena and Glaucoma have higher temperature performance, the prey evolutionary dynamics under Tetrahymena or Glaucoma only treatments show greater similarity to the competition treatment, suggesting a stronger influence of Tetrahymena or Glaucoma in competition treatments under cooler temperatures. Taken together, these results indicate that the temperature performance of different species likely determines the relative dominance among competitors. Furthermore, it appears that the more dominant competitor tends to have a stronger influence on prey evolution.

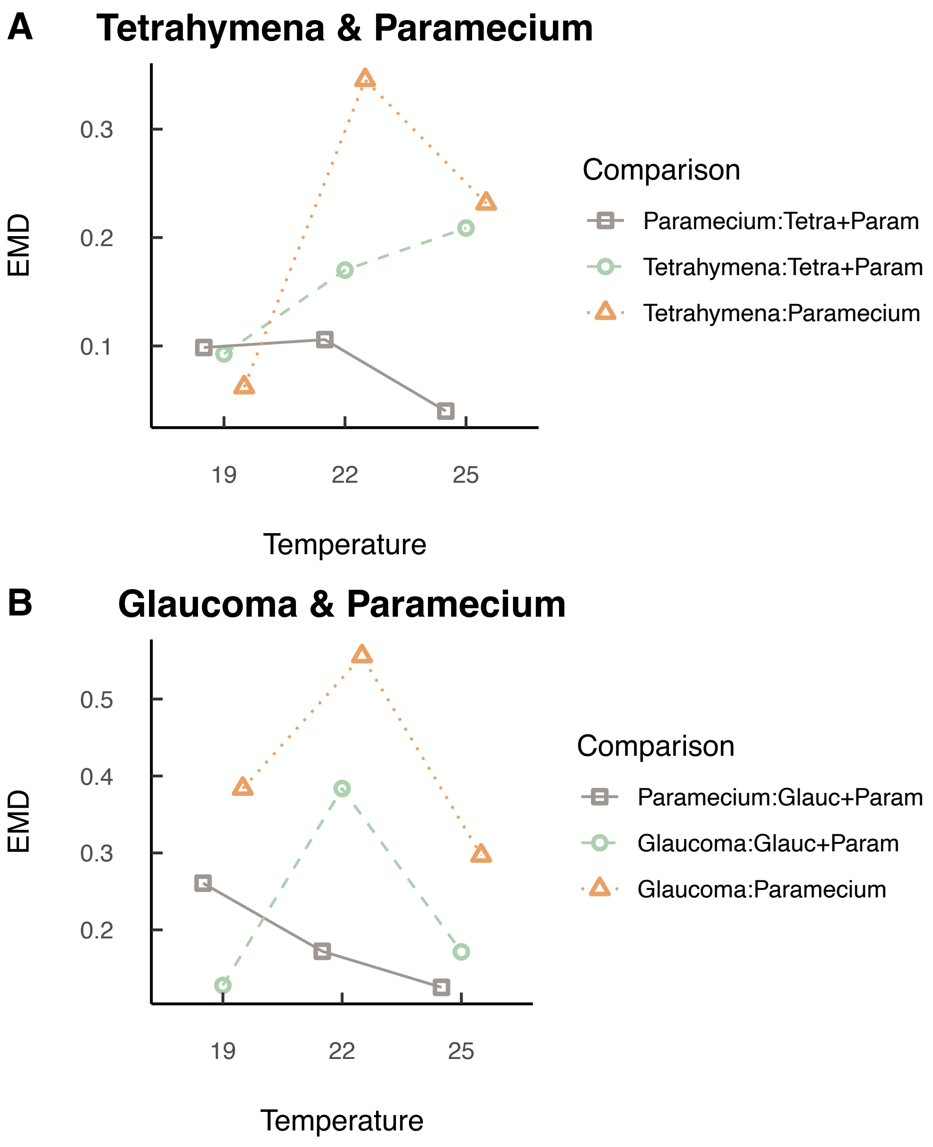


**Figure 1.** Earth Mover's Distance (EMD) of prey evolutionary dynamics (relative frequency of *vfl* overtime) between single predation treatments and competition treatments. Panel A shows the EMDs of pairwise comparison of prey evolutionary dynamics between that of Tetrahymena only, Paramecium only, or Tetrahymena and Paramecium competition treatmtent (Tetra + Param). For instance, solid grey line and open squares represent the EMD between the prey evolutionary dynamics in the Paramecium single predator treatment and the Tetrahymena and Paramecium competition treatment. Similarly, Panel B shows the EMDs of pairwise comparison of Glaucoma only, Paramecium only, or a competitive treatment between the two (Glauc + Param) in Panel B.

### Appendix IV Statistical Analyses

We hypothesized that temperature, predator identity, predator competition, predator density, and time (day) individually and interactively affected prey ecological dynamics (i.e. log density), evolutionary dynamics (i.e. relative frequency of *vfl1-1* and delta frequency of *vfl1-1* across temperature), and prey phenotypic dynamics (change in particle size, measured as log forward scatter height using flow cytometry). We examined our hypotheses using linear mixed models (Table 1-12).

To do so, we first checked for autocorrelation in our data (Table 1). Following this, we compared the single predator treatments (Main text Fig 1b) with control (no predator) to assess? The interactive effects of temperature, predator identity, and day on the ecological dynamics and phenotypic dynamics of WT and *vfl1-1*, and on the relative frequency of *vfl1-1* (Table 2-5,8). We then compared the single predator treatment of *Tetrahymena pyriformis* and *Glaucoma sp.* to two competition treatments. This comparison allowed us to test the four-way interactive effects of temperature, predator identity (*T. pyriformis* and *Glaucoma sp.*), predator competition (presence/absence of *P. caudatum*), and day, on the evolutionary dynamics and phenotypic dynamics of WT and *vfl1-1* (Table 9-12).

To examine how temperature, predator identity, and predator competition affected demographic parameters of both *C. reinhardtii* strain (Table 6) and ach of the protist species (Table 7), we used a linear model to test the population growth rate, *r*, and the maximum population density (proxy for carrying capacity).

For all hypothesis testing (Table 1-5, 8-12), we assumed that the experimental block had random effects on our data while testing the fix effects of temperature, predator identity, predator competition, and day.

**Table 1**. AIC model selection for autoregression with linear mixed models. We performed hypothesis testing on our data to test the main and interactive effects between temperature, predation treatments, and day using 'lmer' in R package 'lme4' (Version 1.1-34). In all models, we treated experimental block as random effects. We tested for autocorrelation using Autoregressive Moving Average (ARMA) correlation structure in the GAMM models using "corARMA" in R package 'nlme' (version 3.1-162). AIC scores indicated that models with no autoregression are the best models (with bolded AIC). Therefore, we used LMM with no autoregressive moving averages for further statistical analysis.

| Response Variables | Models | df | AIC |
|---|---|---|---|
| *wt* log Density (Ecological dynamics) | **log_Density~temperature*predation*day,random = ~ 1\|Block** | 18 | **372.5458** |
| | log_Density~temperature*predation*day,random = ~ 1\|Block, corr = corARMA(form = ~ Day\|Block/Predation/Jar,p=1) | 19 | 374.5458 |
| | log_Density~temperature*predation*day,random = ~ 1\|Block, corr = corARMA(form = ~ Day\|Block/Predation/Jar,p=2) | 20 | 376.5458 |
| | log_Density~temperature*predation*day,random = ~ 1\|Block, corr = corARMA(form = ~ Day\|Block/Predation/Jar,p=3) | 21 | 378.5458 |
| *vfl1-1* log Density (Ecological dynamics) | **log_Density~temperature*predation*day,random = ~ 1\|Block** | 18 | **431.4963** |
| | log_Density~temperature*predation*day,random = ~ 1\|Block, corr = corARMA(form = ~ Day\|Block/Predation/Jar,p=1) | 19 | 433.4963 |
| | log_Density~temperature*predation*day,random = ~ 1\|Block, corr = corARMA(form = ~ Day\|Block/Predation/Jar,p=2) | 20 | 435.4963 |
| | log_Density~temperature*predation*day,random = ~ 1\|Block, corr = corARMA(form = ~ Day\|Block/Predation/Jar,p=3) | 21 | 437.4963 |
| *vfl1-1 relative frequency (Evolutionary dynamics)* | **Frequency~temperature*predation*day,random = ~ 1\|Block** | 18 | **-157.065** |
| | Frequency~temperature*predation*day,random = ~ 1\|Block, corr = corARMA(form = ~ Day\|Block/Predation/Jar,p=1) | 19 | -155.065 |
| | Frequency~temperature*predation*day,random = ~ 1\|Block, corr = corARMA(form = ~ Day\|Block/Predation/Jar,p=2) | 20 | -153.065 |
| | Frequency~temperature*predation*day,random = ~ 1\|Block, corr = corARMA(form = ~ Day\|Block/Predation/Jar,p=3) | 21 | -151.065 |
| *wt size (Phenotypic dynamics)* | **Size~temperature*predation*day,random = ~ 1\|Block** | 18 | **-826.7443** |
| | Size~temperature*predation*day,random = ~ 1\|Block, corr = corARMA(form = ~ Day\|Block/Predation/Jar,p=1) | 19 | -824.7443 |
| | Size~temperature*predation*day,random = ~ 1\|Block, corr = corARMA(form = ~ Day\|Block/Predation/Jar,p=2) | 20 | -822.7443 |
| | Size~temperature*predation*day,random = ~ 1\|Block, corr = corARMA(form = ~ Day\|Block/Predation/Jar,p=3) | 21 | -820.7443 |

**Table 2.** We tested our hypothesis that temperature (Temp), predator identity (single predator treatments, Trt), and time (Day) interactively affected the evolutionary dynamics of *C. reinhardtii* (relative frequency of *vfl1-1*). We compared the relative frequency of *vfl1-1* in single predator treatment (Main Text Fig 1) on days 0, 5, 10, 15 to control (no predator). We tested the individual and interactive effects of listed independent factors using 'lmer' in R package lme4 (version 1.1-34). We assumed that experimental block had random effects in our linear mixed model while the other independent variables had fixed effects. Significance of each independent variable were generated using 'Anova' in package car (version 3.1-2). The p-values for significant independent variables are shown in bold.

| **Frequency ~ Temp * Trt * Day + (1 \| Block)** | | | |
|---|---|---|---|
| **Random effects** | | | |
| Groups | Name | Variance | Std.Dev. |
| Block | (Intercept) | 0.0008205 | 0.02864 |
| Residual | | 0.0144557 | 0.12023 |
| **Fixed effect** | | | |
| | Estimate | Std. Error | t value |
| (Intercept) | 0.2016183 | 0.3917199 | 0.515 |
| Temp | 0.0096216 | 0.0176724 | 0.544 |
| TrtTetra | 0.2021788 | 0.5532346 | 0.365 |
| TrtGlauc | -1.8811831 | 0.5532346 | -3.4 |
| TrtP.cau | -0.8360662 | 0.5532346 | -1.511 |
| Day | -0.0079498 | 0.0362177 | -0.22 |
| Temp:TrtTetra | -0.0169576 | 0.0249926 | -0.679 |
| Temp:TrtGlauc | 0.0848894 | 0.0249926 | 3.397 |
| Temp:TrtP.cau | 0.0486145 | 0.0249926 | 1.945 |
| Temp:Day | -0.0008486 | 0.0016361 | -0.519 |
| TrtTetra:Day | -0.0372834 | 0.0512196 | -0.728 |
| TrtGlauc:Day | 0.2562733 | 0.0512196 | 5.003 |
| TrtP.cau:Day | 0.0985505 | 0.0512196 | 1.924 |
| Temp:TrtTetra:Day | 0.003387 | 0.0023139 | 1.464 |
| Temp:TrtGlauc:Day | -0.0096477 | 0.0023139 | -4.17 |
| Temp:TrtP.cau:Day | -0.0051354 | 0.0023139 | -2.219 |
| **Significance** | | | |
| Variable | Chisq | Df | Pr(>Chisq) |
| Temp | 0.2888 | 1 | 0.59098 |
| Trt | 369.7689 | 3 | **< 2.2e-16** |
| Day | 30.0803 | 1 | **4.15E-08** |
| Temp:Trt | 9.242 | 3 | **0.02624** |
| Temp:Day | 19.7166 | 1 | **8.98E-06** |
| Trt:Day | 148.6746 | 3 | **< 2.2e-16** |
| Temp:Trt:Day | 36.07 | 3 | **7.24E-08** |

**Table 3.** Hypothesis testing for the interactive effects between temperature (Temp), single predator treatments (Trt), and time (Day) on the phenotypic dynamics of *C. reinhardtii vfl1-1* mutant strain (mean log forward scatter height).

| **MeanClumpSize~Temp * Trt * Day + (1 \| Block)** | | | |
|---|---|---|---|
| **Random effects** | | | |
| Groups | Name | Variance | Std.Dev. |
| Block | (Intercept) | 0.0001722 | 0.01312 |
| Residual | | 0.0016197 | 0.04025 |
| **Fixed effect** | | | |
| | Estimate | Std. Error | t value |
| (Intercept) | 13.99 | 0.07232 | 193.415 |
| Temp | -0.001388 | 0.00324 | -0.428 |
| TrtTetra | 0.04849 | 0.1014 | 0.478 |
| TrtGlauc | 0.09549 | 0.1014 | 0.941 |
| TrtP.cau | -0.0393 | 0.1014 | -0.387 |
| Day | 0.0085 | 0.007667 | 1.109 |
| Temp:TrtTetra | -0.00242 | 0.004582 | -0.528 |
| Temp:TrtGlauc | -0.00405 | 0.004582 | -0.884 |
| Temp:TrtP.cau | 0.001114 | 0.004582 | 0.243 |
| Temp:Day | -0.0003755 | 0.0003464 | -1.084 |
| TrtTetra:Day | -0.0107 | 0.01084 | -0.987 |
| TrtGlauc:Day | 0.0002475 | 0.01084 | 0.023 |
| TrtP.cau:Day | 0.000402 | 0.01084 | 0.037 |
| Temp:TrtTetra:Day | 0.0006528 | 0.0004899 | 1.333 |
| Temp:TrtGlauc:Day | 0.0001099 | 0.0004899 | 0.224 |
| Temp:TrtP.cau:Day | 0.00005096 | 0.0004899 | 0.104 |
| **Significance** | | | |
| Variable | Chisq | Df | Pr(>Chisq) |
| Temp | 17.2225 | 1 | **0.00003325** |
| Trt | 31.1177 | 3 | **8.029E-07** |
| Day | 26.8911 | 1 | **2.15E-07** |
| Temp:Trt | 4.9715 | 3 | 0.1739 |
| Temp:Day | 0.9879 | 1 | 3.20E-01 |
| Trt:Day | 10.3122 | 3 | **0.01609** |
| Temp:Trt:Day | 2.2947 | 3 | 5.14E-01 |

**Table 4.** Hypothesis testing for interactive effects between temperature (Temp), single predator treatments (Trt), and time (Day) on the phenotypic dynamics of *C. reinhardtii* wild type (mean log forward scatter height). Methods details see Table 2. Both versions of the models show cell clump size of WT was significantly affected by all independent effects and all two-way interactive effects of temperature, predator species identity, and time.

| **MeanClumpSize~ Temp * Trt * Day + (1 \| Block)** | | | |
|---|---|---|---|
| **Random effects** | | | |
| Groups | Name | Variance | Std.Dev. |
| Block | (Intercept) | 0.0006423 | 0.02534 |
| Residual | | 0.0115284 | 0.10737 |
| **Fixed effect** | | | |
| | Estimate | Std. Error | t value |
| (Intercept) | 13.9914305 | 0.1921832 | 72.803 |
| Temp | -0.0028179 | 0.0086441 | -0.326 |
| TrtTetra | -0.2443791 | 0.270604 | -0.903 |
| TrtGlauc | -0.3074343 | 0.270604 | -1.136 |
| TrtP.cau | -0.6088699 | 0.270604 | -2.25 |
| Day | -0.003583 | 0.0204557 | -0.175 |
| Temp:TrtTetra | 0.0106773 | 0.0122246 | 0.873 |
| Temp:TrtGlauc | 0.0140716 | 0.0122246 | 1.151 |
| Temp:TrtP.cau | 0.0254439 | 0.0122246 | 2.081 |
| Temp:Day | 0.0004659 | 0.0009241 | 0.504 |
| TrtTetra:Day | 0.0228992 | 0.0289288 | 0.792 |
| TrtGlauc:Day | 0.0692475 | 0.0289288 | 2.394 |
| TrtP.cau:Day | -0.0006055 | 0.0289288 | -0.021 |
| Temp:TrtTetra:Day | -0.0007906 | 0.0013069 | -0.605 |
| Temp:TrtGlauc:Day | -0.0023937 | 0.0013069 | -1.832 |
| Temp:TrtP.cau:Day | 0.0005687 | 0.0013069 | 0.435 |
| **Significance** | | | |
| Variable | Chisq | Df | Pr(>Chisq) |
| Variable | Chisq | Df | Pr(>Chisq) |
| Temp | 10.3776 | 1 | **0.001276** |
| Trt | 54.8689 | 3 | **7.323E-12** |
| Day | 179.5952 | 1 | **< 2.2e-16** |
| Temp:Trt | 25.7255 | 3 | **1.09E-05** |
| Temp:Day | 0.1655 | 1 | 0.684175 |
| Trt:Day | 30.8786 | 3 | **9.016E-07** |
| Temp:Trt:Day | 5.8174 | 3 | 0.120841 |

**Table 5.** Temperature and predator competition effects on *C. reinhardtii* growth rate, *r*, and maximum population.

| **r ~ Temp * Trt * Strain** | | | | |
|---|---|---|---|---|
| Estimate | Estimate | Std. Error | t value | Pr(>\|t\|) |
| (Intercept) | -0.515115 | 0.178985 | -2.878 | **0.004466** |
| Temp | 0.039973 | 0.008086 | 4.944 | **1.69E-06** |
| TrtTetra | 0.474472 | 0.253123 | 1.874 | 0.062417 |
| TrtGlauc | 0.391925 | 0.253123 | 1.548 | 0.123218 |
| TrtP.cau | 0.761374 | 0.253123 | 3.008 | **0.002991** |
| TrtTetraP.cau | 1.151157 | 0.260363 | 4.421 | **1.66E-05** |
| TrtGlaucP.cau | 0.719444 | 0.253123 | 2.842 | **0.004974** |
| StrainWT-mNG | 0.326109 | 0.253123 | 1.288 | 0.19921 |
| Temp:TrtTetra | -0.011576 | 0.011435 | -1.012 | 0.312663 |
| Temp:TrtGlauc | -0.00679 | 0.011435 | -0.594 | 0.553372 |
| Temp:TrtP.cau | -0.026842 | 0.011435 | -2.347 | **0.019946** |
| Temp:TrtTetraP.cau | -0.039324 | 0.011709 | -3.358 | **0.000949** |
| Temp:TrtGlaucP.cau | -0.016146 | 0.011435 | -1.412 | 0.159604 |
| Temp:StrainWT-mNG | -0.006761 | 0.011435 | -0.591 | 0.555058 |
| TrtTetra:StrainWT-mNG | 0.160305 | 0.35797 | 0.448 | 0.654802 |
| TrtGlauc:StrainWT-mNG | 0.283191 | 0.35797 | 0.791 | 0.429882 |
| TrtP.cau:StrainWT-mNG | 0.573099 | 0.35797 | 1.601 | 0.111063 |
| TrtTetraP.cau:StrainWT-mNG | 0.929549 | 0.368208 | 2.525 | **0.012413** |
| TrtGlaucP.cau:StrainWT-mNG | 0.345889 | 0.35797 | 0.966 | 0.33516 |
| Temp:TrtTetra:StrainWT-mNG | -0.007161 | 0.016171 | -0.443 | 0.658422 |
| Temp:TrtGlauc:StrainWT-mNG | -0.01791 | 0.016171 | -1.107 | 0.269501 |
| Temp:TrtP.cau:StrainWT-mNG | -0.031942 | 0.016171 | -1.975 | **0.049709** |
| Temp:TrtTetraP.cau:StrainWT-mNG | -0.048673 | 0.01656 | -2.939 | **0.003702** |
| Temp:TrtGlaucP.cau:StrainWT-mNG | -0.025249 | 0.016171 | -1.561 | 0.120121 |
| Adjusted R-squared: 0.6748 ; model p-value: < 2.2e-16 | | | | |
| **Maximum_density ~ Temp * Trt * Strain** | | | | |
| Estimate | Estimate | Std. Error | t value | Pr(>\|t\|) |
| (Intercept) | 6.075906 | 0.946289 | 6.421 | **1.08E-09** |
| Temp | 0.156092 | 0.042749 | 3.651 | **3.38E-04** |
| TrtTetra | 1.539057 | 1.338255 | 1.15 | 0.251585 |
| TrtGlauc | 4.690177 | 1.338255 | 3.505 | **0.000571** |
| TrtP.cau | 2.756387 | 1.338255 | 2.06 | **0.040806** |
| TrtTetraP.cau | 4.705223 | 1.376529 | 3.418 | **7.73E-04** |
| TrtGlaucP.cau | 2.963145 | 1.338255 | 2.214 | **0.028018** |
| StrainWT-mNG | 4.136753 | 1.338255 | 3.091 | **0.002297** |
| Temp:TrtTetra | -0.012282 | 0.060456 | -0.203 | 0.839239 |
| Temp:TrtGlauc | -0.142158 | 0.060456 | -2.351 | **0.019737** |
| Temp:TrtP.cau | -0.090442 | 0.060456 | -1.496 | 0.136334 |
| Temp:TrtTetraP.cau | -0.152846 | 0.061907 | -2.469 | **0.014443** |
| Temp:TrtGlaucP.cau | -0.054313 | 0.060456 | -0.898 | 0.370131 |
| Temp:StrainWT-mNG | -0.094494 | 0.060456 | -1.563 | 0.119732 |
| TrtTetra:StrainWT-mNG | -1.480679 | 1.892579 | -0.782 | 0.434987 |
| TrtGlauc:StrainWT-mNG | -4.871346 | 1.892579 | -2.574 | **0.010825** |
| TrtP.cau:StrainWT-mNG | -4.042957 | 1.892579 | -2.136 | **0.033955** |
| TrtTetraP.cau:StrainWT-mNG | -0.112408 | 1.946706 | -0.058 | 0.954015 |
| TrtGlaucP.cau:StrainWT-mNG | -1.193267 | 1.892579 | -0.63 | 0.529134 |
| Temp:TrtTetra:StrainWT-mNG | 0.005957 | 0.085498 | 0.07 | 0.94453 |
| Temp:TrtGlauc:StrainWT-mNG | 0.123121 | 0.085498 | 1.44 | 0.151518 |
| Temp:TrtP.cau:StrainWT-mNG | 0.123501 | 0.085498 | 1.444 | 0.150266 |
| Temp:TrtTetraP.cau:StrainWT-mNG | -0.069979 | 0.08755 | -0.799 | 0.425125 |
| Temp:TrtGlaucP.cau:StrainWT-mNG | -0.048204 | 0.085498 | -0.564 | 0.573563 |
| Adjusted R-squared: 0.6219 ; model p-value: < 2.2e-16 |  |  |  |  |

**Table 6.** Hypothesis testing for interactive effects between temperature (Temp), single predator treatments (Trt), and time (Day) on the Δ *vfl1-1* frequency across temperature. All main effects and interactive effects of the independent variables are significant, further indicating that temperature effects are mediated through predator-prey interactions.

| **Delta_Frequency ~ Temp * Trt * Day + (1 \| Block)** | | | |
|---|---|---|---|
| **Random effects** | | | |
| Groups | Name | Variance | Std.Dev. |
| Block | (Intercept) | 0.0006408 | 0.02531 |
| Residual | | 0.0170045 | 0.1304 |
| **Fixed effect** | | | |
| | Estimate | Std. Error | t value |
| (Intercept) | -0.1117862 | 0.9028676 | -0.124 |
| Temp | 0.0066879 | 0.0383343 | 0.174 |
| TrtTetra | 0.583562 | 1.2765967 | 0.457 |
| TrtGlauc | -1.2982234 | 1.2765967 | -1.017 |
| TrtP.cau | -1.1776216 | 1.2765967 | -0.922 |
| Day | -0.0088638 | 0.0835729 | -0.106 |
| Temp:TrtTetra | -0.0273105 | 0.0542129 | -0.504 |
| Temp:TrtGlauc | 0.0714513 | 0.0542129 | 1.318 |
| Temp:TrtP.cau | 0.0578508 | 0.0542129 | 1.067 |
| Temp:Day | 0.0001529 | 0.0035491 | 0.043 |
| TrtTetra:Day | 0.0538845 | 0.1181899 | 0.456 |
| TrtGlauc:Day | 0.2851861 | 0.1181899 | 2.413 |
| TrtP.cau:Day | 0.0971851 | 0.1181899 | 0.822 |
| Temp:TrtTetra:Day | -0.0013615 | 0.0050191 | -0.271 |
| Temp:TrtGlauc:Day | -0.0136018 | 0.0050191 | -2.71 |
| Temp:TrtP.cau:Day | -0.0050314 | 0.0050191 | -1.002 |
| **Significance** | | | |
| Variable | Chisq | Df | Pr(>Chisq) |
| Temp | 5.0449 | 1 | **0.024698** |
| Trt | 44.1835 | 3 | **1.38E-09** |
| Day | 26.3947 | 1 | **2.78E-07** |
| Temp:Trt | 16.6805 | 3 | **0.0008221** |
| Temp:Day | 7.4568 | 1 | **0.0063196** |
| Trt:Day | 64.4692 | 3 | **6.52E-14** |
| Temp:Trt:Day | 8.9102 | 3 | **0.0305094** |

**Table 7.** Temperature and predator competition effects on predator population growth rate, *r*, and maximum population. Models show that temperature had significant independent and interactive on predator population growth rate, but only significant interactive effects on predator maximum population density. The *r* and maximum population growth rate of *Glaucoma sp.* did not differ significantly from that of *T. pyriformis* (intercept) in the absence of competition, but did in competition with *P. caudatum*. This result further supports that *Glaucoma sp.* and *T. pyriformis*, despite having similar size and shape, differ in many aspects including their competitive abilities with *P. caudatum*.

| **r ~ Temp + Species + PredComp + Temp:Species + Species:PredComp** | | | | |
|---|---|---|---|---|
| Estimate | Estimate | Std. Error | t value | Pr(>\|t\|) |
| (Intercept) | 1.437804 | 0.140004 | 10.27 | **< 2e-16** |
| Temp | -0.014927 | 0.00629 | -2.373 | **0.01939** |
| SpeciesGlauc | 0.09105 | 0.195844 | 0.465 | 0.64292 |
| SpeciesP.cau | -3.454961 | 0.186048 | -18.57 | **< 2e-16** |
| PredCompYes | -0.145236 | 0.031073 | -4.674 | **8.50E-06** |
| Temp:SpeciesGlauc | -0.012012 | 0.008796 | -1.366 | 0.17487 |
| Temp:SpeciesP.cau | 0.138482 | 0.008345 | 16.594 | **< 2e-16** |
| SpeciesGlauc:PredCompYes | 0.218383 | 0.043278 | 5.046 | **1.82E-06** |
| SpeciesP.cau:PredCompYes | 0.125427 | 0.041219 | 3.043 | **0.00294** |
| Adjusted R-squared: 0.877 ; model p-value: < 2.2e-16 | | | | |
| **Maximum_density ~ Temp + Species + PredComp + Temp:Species + Species:PredComp** | | | | |
| Estimate | Estimate | Std. Error | t value | Pr(>\|t\|) |
| (Intercept) | 8.50287 | 0.65594 | 12.963 | **< 2e-16** |
| Temp | -0.04867 | 0.02947 | -1.652 | 0.10138 |
| SpeciesGlauc | 1.26327 | 0.91756 | 1.377 | 0.1713 |
| SpeciesP.cau | -6.38889 | 0.84611 | -7.551 | **1.20E-11** |
| PredCompYes | -0.882 | 0.14558 | -6.058 | **1.85E-08** |
| Temp:SpeciesGlauc | -0.07214 | 0.04121 | -1.751 | 0.08273 |
| Temp:SpeciesP.cau | 0.17869 | 0.03792 | 4.712 | **7.03E-06** |
| SpeciesGlauc:PredCompYes | 0.52456 | 0.20276 | 2.587 | **0.01095** |
| SpeciesP.cau:PredCompYes | 0.63571 | 0.19087 | 3.331 | **0.00117** |
| Adjusted R-squared: 0.8765 ; model p-value: < 2.2e-16 | | | | |

**Table 8.** We tested how temperature, predator treatment, and day individually and interactively affect the median, variance, skewness and kurtosis of WT particle size distribution. We only present the p-value for the model terms.

| | Mediam | SD | Skewness | Kurtosis |
|---|---|---|---|---|
| **Temp** | **0.02049** | 0.48634 | **1.08E-08** | **1.16E-08** |
| **Trt** | **0.00008765** | **< 2e-16** | **< 2.2e-16** | **9.10E-12** |
| **Day** | **< 2.2e-16** | **< 2e-16** | **< 2.2e-16** | **6.62E-06** |
| **Temp:Trt** | **4.44E-06** | **0.04705** | **3.90E-05** | **0.0001986** |
| **Temp:Day** | 0.69015 | 0.15568 | 0.20542 | **0.0195878** |
| **Trt:Day** | **0.00356** | **< 2e-16** | **1.08E-08** | **4.88E-12** |
| **Temp:Trt:Day** | **0.04659** | 0.08892 | **0.03326** | 0.2677402 |

**Table 9.** Hypothesis testing for interactive effects between temperature, predator identity (*T. pyriformis* and *Glaucoma sp.*), predator competition (presence/absence of *P. caudatum*), and day on the evolutionary dynamics of *vfl1-1*. Model shows that four-way interaction is significant.

| **vfl_Freq ~ Temp * FocalSp * PredComp * Day + (1 \| Block)** | | | |
|---|---|---|---|
| **Random effects** | | | |
| Groups | Name | Variance | Std.Dev. |
| Block | (Intercept) | 0.001802 | 0.04245 |
| Residual | | 0.018286 | 0.13522 |
| **Fixed effect** | | | |
| | Estimate | Std. Error | t value |
| (Intercept) | -1.596158 | 0.440999 | -3.619 |
| Temp | 0.090578 | 0.019876 | 4.557 |
| FocalSpTetra | 2.006059 | 0.622221 | 3.224 |
| PredCompYes | 1.012099 | 0.622221 | 1.627 |
| Day | 0.234936 | 0.040734 | 5.768 |
| Temp:FocalSpTetra | -0.097877 | 0.028109 | -3.482 |
| Temp:PredCompYes | -0.031308 | 0.028109 | -1.114 |
| FocalSpTetra:PredCompYes | -2.661358 | 0.89266 | -2.981 |
| Temp:Day | -0.009983 | 0.00184 | -5.425 |
| FocalSpTetra:Day | -0.279911 | 0.057606 | -4.859 |
| PredCompYes:Day | -0.090893 | 0.057606 | -1.578 |
| Temp:FocalSpTetra:PredCompYes | 0.124739 | 0.040233 | 3.1 |
| Temp:FocalSpTetra:Day | 0.012451 | 0.002602 | 4.785 |
| Temp:PredCompYes:Day | 0.001991 | 0.002602 | 0.765 |
| FocalSpTetra:PredCompYes:Day | 0.318247 | 0.082641 | 3.851 |
| Temp:FocalSpTetra:PredCompYes:Da | -0.01427 | 0.003725 | -3.831 |
| **Significance** | | | |
| Variable | Chisq | Df | Pr(>Chisq) |
| Temp | 2.4883 | 1 | 0.1146936 |
| FocalSp | 60.7623 | 1 | **6.44E-15** |
| PredComp | 22.1872 | 1 | **2.473E-06** |
| Day | 18.8159 | 1 | **1.44E-05** |
| Temp:FocalSp | 5.5249 | 1 | **0.0187479** |
| Temp:PredComp | 7.0335 | 1 | **0.0080001** |
| FocalSp:PredComp | 11.1549 | 1 | **0.0008381** |
| Temp:Day | 45.136 | 1 | **1.84E-11** |
| FocalSp:Day | 0.7595 | 1 | 0.3834982 |
| PredComp:Day | 99.21 | 1 | **< 2.2e-16** |
| Temp:FocalSp:PredComp | 1.3952 | 1 | 0.2375304 |
| Temp:FocalSp:Day | 8.6794 | 1 | **0.0032183** |
| Temp:PredComp:Day | 7.1394 | 1 | **0.0075407** |
| FocalSp:PredComp:Day | 0.1529 | 1 | 0.6957689 |
| Temp:FocalSp:PredComp:Day | 14.6783 | 1 | **1.28E-04** |

**Table 10.** Hypothesis testing for interactive effects between predator identity (*T. pyriformis* and *Glaucoma sp.*), predator competition (presence/absence of *P. caudatum*), and day on the delta changes in relative frequency of *vfl1-1* across temperature (using its relative frequency at 19℃ as the baseline). The model indicates that predator identity and predator competition significantly affect the temperature effects on *vfl1-1* relative frequency. This further supports the hypothesis that temperature effects on *C. reinhardtii* evolution is most likely mediated through biotic interactions.

| Delta_Freq ~ Temp * FocalSp * PredComp * Day + (1 \| Block) | | | |
|---|---|---|---|
| **Random effects** | | | |
| Groups | Name | Variance | Std.Dev. |
| Block | (Intercept) | 0.001345 | 0.03667 |
| Residual | | 0.022182 | 0.14894 |
| **Fixed effect** | | | |
| | Estimate | Std. Error | t value |
| (Intercept) | -1.41001 | 1.03132 | -1.367 |
| Temp | 0.078139 | 0.043783 | 1.785 |
| FocalSpTetra | 1.881785 | 1.458046 | 1.291 |
| PredCompYes | 1.508291 | 1.458046 | 1.034 |
| Day | 0.276322 | 0.095452 | 2.895 |
| Temp:FocalSpTetra | -0.098762 | 0.061919 | -1.595 |
| Temp:PredCompYes | -0.067846 | 0.061919 | -1.096 |
| FocalSpTetra:PredCompYes | -1.533108 | 2.090924 | -0.733 |
| Temp:Day | -0.013449 | 0.004054 | -3.318 |
| FocalSpTetra:Day | -0.231302 | 0.134989 | -1.713 |
| PredCompYes:Day | -0.285098 | 0.134989 | -2.112 |
| Temp:FocalSpTetra:PredCompYes | 0.092076 | 0.088657 | 1.039 |
| Temp:FocalSpTetra:Day | 0.01224 | 0.005733 | 2.135 |
| Temp:PredCompYes:Day | 0.011882 | 0.005733 | 2.073 |
| FocalSpTetra:PredCompYes:Day | 0.24021 | 0.193574 | 1.241 |
| Temp:FocalSpTetra:PredCompYes:Da | -0.013086 | 0.008208 | -1.594 |
| **Significance** | | | |
| Variable | Chisq | Df | Pr(>Chisq) |
| Temp | 11.9439 | 1 | **5.48E-04** |
| FocalSp | 16.8695 | 1 | **4.00E-05** |
| PredComp | 43.194 | 1 | **4.96E-11** |
| Day | 100.8705 | 1 | **< 2.2e-16** |
| Temp:FocalSp | 0.0799 | 1 | 0.7774718 |
| Temp:PredComp | 3.6634 | 1 | 0.0556203 |
| FocalSp:PredComp | 0.7584 | 1 | 0.3838313 |
| Temp:Day | 5.2811 | 1 | **0.0215581** |
| FocalSp:Day | 14.5362 | 1 | **0.0001375** |
| PredComp:Day | 39.7644 | 1 | **2.87E-10** |
| Temp:FocalSp:PredComp | 1.3387 | 1 | 0.2472641 |
| Temp:FocalSp:Day | 2.0382 | 1 | 0.153391 |
| Temp:PredComp:Day | 1.7963 | 1 | 0.1801606 |
| FocalSp:PredComp:Day | 30.3259 | 1 | **3.65E-08** |
| Temp:FocalSp:PredComp:Day | 2.5418 | 1 | 0.1108713 |

**Table 11.** Hypothesis testing for interactive effects between temperature, predator identity (*T. pyriformis* and *Glaucoma sp.*), predator competition (presence/absence of *P. caudatum*), and day on the phenotypic dynamics of WT.

| MeanClumpSize ~ Temp * FocalSp * PredComp * Day + (1 \| Block) | | | |
|---|---|---|---|
| **Random effects** | | | |
| Groups | Name | Variance | Std.Dev. |
| Block | (Intercept) | 0.001242 | 0.03524 |
| Residual | | 0.020987 | 0.14487 |
| **Fixed effect** | | | |
| | Estimate | Std. Error | t value |
| (Intercept) | 13.68 | 0.2594 | 52.759 |
| Temp | 0.01125 | 0.01166 | 0.965 |
| FocalSpTetra | 0.06306 | 0.3651 | 0.173 |
| PredCompYes | -0.5079 | 0.3651 | -1.391 |
| Day | 0.06566 | 0.0276 | 2.379 |
| Temp:FocalSpTetra | -0.003394 | 0.01649 | -0.206 |
| Temp:PredCompYes | 0.02356 | 0.01649 | 1.428 |
| FocalSpTetra:PredCompYes | -0.2918 | 0.5238 | -0.557 |
| Temp:Day | -0.001928 | 0.001247 | -1.546 |
| FocalSpTetra:Day | -0.04635 | 0.03903 | -1.187 |
| PredCompYes:Day | 0.003306 | 0.03903 | 0.085 |
| Temp:FocalSpTetra:PredCompYes | 0.01315 | 0.02361 | 0.557 |
| Temp:FocalSpTetra:Day | 0.001603 | 0.001763 | 0.909 |
| Temp:PredCompYes:Day | 0.00008813 | 0.001763 | 0.05 |
| FocalSpTetra:PredCompYes:Day | 0.04815 | 0.05599 | 0.86 |
| Temp:FocalSpTetra:PredCompYes:Day | -0.001546 | 0.002524 | -0.612 |
| **Significance** | | | |
| Variable | Chisq | Df | Pr(>Chisq) |
| Temp | 14.0306 | 1 | **1.80E-04** |
| FocalSp | 6.4687 | 1 | **0.0109789** |
| PredComp | 33.9756 | 1 | **5.58E-09** |
| Day | 231.7476 | 1 | **< 2.2e-16** |
| Temp:FocalSp | 1.7713 | 1 | 0.1832258 |
| Temp:PredComp | 12.5381 | 1 | **0.0003987** |
| FocalSp:PredComp | 8.9114 | 1 | **0.0028339** |
| Temp:Day | 5.374 | 1 | **2.04E-02** |
| FocalSp:Day | 1.8569 | 1 | 0.1729866 |
| PredComp:Day | 15.0421 | 1 | **0.0001051** |
| Temp:FocalSp:PredComp | 0.0121 | 1 | 0.9122751 |
| Temp:FocalSp:Day | 0.4524 | 1 | 0.5011773 |
| Temp:PredComp:Day | 0.2791 | 1 | 0.597315 |
| FocalSp:PredComp:Day | 5.1359 | 1 | **0.0234361** |
| Temp:FocalSp:PredComp:Day | 0.3751 | 1 | 0.5402228 |

**Table 12.** Hypothesis testing for interactive effects between temperature, predator identity (*T. pyriformis* and *Glaucoma sp.*), predator competition (presence/absence of *P. caudatum*), and day on the phenotypic dynamics of *vfl1-1*.

| **MeanClumpSize ~ Temp * FocalSp * PredComp * Day + (1 \| Block)** | | | |
|---|---|---|---|
| **Random effects** | | | |
| Groups | Name | Variance | Std.Dev. |
| Block | (Intercept) | 0.0001164 | 0.01079 |
| Residual | | 0.0011988 | 0.03462 |
| **Fixed effect** | | | |
| | Estimate | Std. Error | t value |
| (Intercept) | 14.0833734 | 0.0621727 | 226.52 |
| Temp | -0.0054382 | 0.0027874 | -1.951 |
| FocalSpTetra | -0.0469954 | 0.0872609 | -0.539 |
| PredCompYes | -0.1574904 | 0.0872609 | -1.805 |
| Day | 0.0087471 | 0.0065963 | 1.326 |
| Temp:FocalSpTetra | 0.0016304 | 0.003942 | 0.414 |
| Temp:PredCompYes | 0.0065168 | 0.003942 | 1.653 |
| FocalSpTetra:PredCompYes | 0.1139954 | 0.1251952 | 0.911 |
| Temp:Day | -0.0002657 | 0.000298 | -0.892 |
| FocalSpTetra:Day | -0.0109476 | 0.0093286 | -1.174 |
| PredCompYes:Day | 0.0170876 | 0.0093286 | 1.832 |
| Temp:FocalSpTetra:PredCompYes | -0.0050712 | 0.0056426 | -0.899 |
| Temp:FocalSpTetra:Day | 0.0005429 | 0.0004214 | 1.288 |
| Temp:PredCompYes:Day | -0.0008053 | 0.0004214 | -1.911 |
| FocalSpTetra:PredCompYes:Day | -0.0107935 | 0.0133826 | -0.807 |
| Temp:FocalSpTetra:PredCompYes:Day | 0.0005449 | 0.0006032 | 0.903 |
| **Significance** | | | |
| Variable | Chisq | Df | Pr(>Chisq) |
| Temp | 30.3527 | 1 | **3.60E-08** |
| FocalSp | 0.2549 | 1 | 0.613648 |
| PredComp | 10.377 | 1 | **0.001276** |
| Day | 82.1341 | 1 | **< 2.2e-16** |
| Temp:FocalSp | 9.5958 | 1 | **0.00195** |
| Temp:PredComp | 0 | 1 | 0.998421 |
| FocalSp:PredComp | 1.8676 | 1 | 0.171746 |
| Temp:Day | 3.1384 | 1 | 0.076469 |
| FocalSp:Day | 4.8205 | 1 | **0.028124** |
| PredComp:Day | 0.0072 | 1 | 0.93236 |
| Temp:FocalSp:PredComp | 0.0852 | 1 | 0.770387 |
| Temp:FocalSp:Day | 7.1984 | 1 | **0.007297** |
| Temp:PredComp:Day | 3.1997 | 1 | 7.37E-02 |
| FocalSp:PredComp:Day | 0.6787 | 1 | 0.410024 |
| Temp:FocalSp:PredComp:Day | 0.8162 | 1 | 0.366305 |
